# ARGON: Fast, Whole-Genome Simulation of the Discrete Time Wright-Fisher Process

**DOI:** 10.1101/036376

**Authors:** Pier Francesco Palamara

## Abstract

**Motivation:** Simulation under the coalescent model is ubiquitous in the analysis of genetic data. The rapid growth of real data sets from multiple human populations led to increasing interest in simulating very large sample sizes at whole-chromosome scales. When the sample size is large, the coalescent model becomes an increasingly inaccurate approximation of the discrete time Wright-Fisher model (DTWF). Analytical and computational treatment of the DTWF, however, is generally harder.

**Results:** We present a simulator (ARGON) for the DTWF process that scales up to hundreds of thousands of samples and whole-chromosome lengths, with a time/memory performance comparable or superior to currently available methods for coalescent simulation. The simulator supports arbitrary demographic history, migration, Newick tree output, variable mutation/recombination rates and gene conversion, and efficiently outputs pairwise identical-by-descent (IBD) sharing data.

**Availability:** ARGON (version 0.1) is written in Java, open source, and freely available at https://github.com/pierpal/ARGON.

**Supplementary information:** Supplementary data are available online.

## 1 Introduction

The coalescent ([Kingman, 1982]) can be constructed as an approximation of the discrete time Wright-Fisher process (DTWF, [Fisher et al., 1922, Wright, 1931]), and leads to simplified analytical and computational treatment. Simulators based on the coalescent process (e.g. [Hudson, 2002]) have been extensively adopted in computational methods. The coalescent approximation, however, relies on the assumption that the sample size is small compared to the effective population size, and violations of this assumption may result in substantial distortions of key genealogical properties ([Wakeley and Takahashi, 2003, Bhaskar et al., 2014]).

Until very recently, coalescent simulators did not scale up to long chromosomes and the very large sample sizes of modern day data sets, which now comprise hundreds of thousands of individuals. The recently developed simulators COSI2 ([Shlyakhter et al., 2014]) and SCRM ([Staab et al., 2015]) enable fast simulation under approximate coalescent models. The COSI algorithm uses a standard backwards-in-time approach, while SCRM adopts a “sequential” approach ([Wiuf and Hein, 2000, McVean and Cardin, 2005]). Both can be also used to simulate large sample sizes and chromosome-long regions under the coalescent process with reasonable time and memory requirements.

Here, we present ARGON, an efficient simulator of the DTWF process that scales up to very large chromosomes, and hundreds of thoudsands of samples. The simulator offers substantially improved performance compared to recent DTWF simulators, e.g. GENOME ([Liang et al., 2007]), and is comparable or superior to current coalescent simulators in terms of speed and memory usage.

## 2 Approach

ARGON proceeds backwards in time one generation at a time, occasionally sampling coalescent and recombination events subject to population structure and migration. Each individual is represented as a list of regions that are still being tracked at the current time (i.e. for which not all samples have found a common ancestor). Recombination events are sampled in genetic space from an exponential distribution, and rounded to the closest physiscal base pair position based on the desired recombination rate, which may be varying along the chromosome. A recombination event can be a crossover or a non-crossover event, based on user-specified rates. For each individual, a maximum of two parents are sampled, so that multiple recombination events result in alternating ancestry between two individuals from the previous generation. When two or more individuals choose the same parent, coalescence occurs if the individuals contain overlapping regions of genetic material. During coalescence, regions within individuals are annotated with links to descendant ancestral recombination graph (ARG [Griffiths, 1981]) nodes.

Compared to other DTWF implementations, ARGON offers substantially improved speed and memory usage. In the GENOME simulator, for instance, individuals are represented as arrays of blocks of genetic material of a fixed size, and individuals from all populations are explicitly represented in memory using arrays. In ARGON, large regions are represented as intervals with arbitrary boundary values, and hash map data structures are extensively used to take advantage of sparsity, avoiding explicit representation of all individuals. As in the GENOME simulator, ARGON can run in approximate mode, so that recombinations breakpoints are rounded to blocks of a user-specified genetic length. This reduces the granularity of the recombination process, improving speed and memory usage, at the cost of slightly inflated correlation of markers at a short genomic distance. We tested this approximation using nonrecombinant blocks of size 10*μM* (1*μM* = 0.000001 Morgans). Additional tests using 50*μM* blocks are described in the Supplementary Note.

ARGON can efficiently output a list of pairwise IBD segments longer than a user-specified centimorgan length. Previous approaches to output simulated IBD sharing data (e.g. [Palamara et al., 2012, Palamara and Pe’er, 2013, Browning and Browning, 2013]) required comparing recent ancestry for all pairs of individual at each marginal tree in the sampled ARG, with computational cost quadratic in the sample size and linear in the number of analyzed marginal trees. IBD segments are delimited by the occurrence of recombination events that change the most recent common ancestor for pairs of samples. In ARGON, these events are detected by visiting internal ARG nodes, which requires less computational effort than the approach based on marginal trees.

## 3 Results

### 3.1 Accuracy for small sample sizes

When *n ≪ N_e_*, the coalescent is a good approximation of the DTWF. We performed extensive testing for several scenarios including population size variation, migration across multiple demes, and gene conversion. We report detailed results in the Supplementary Note. We find good agreement between ARGON and MS. We also tested the accuracy of COSI version 2.0, SCRM version 1.6.1, and MSprime version 0.1.6 ([Kelleher et al., 2015]), a new efficient simulator for which a preliminary version was released at the time of writing. All simulators were found to be well calibrated against MS.

**Table 1:** Comparison of theoretical predictions and simulation results for the number of singleton and doubleton alleles when *n* = *N_e_*. We performed 100, 000 independent simulations, for a region of 1 Mb, *μ* = 2 × 10^−8^, and 1 cM/Mb. Reported errors are obtained as 100 × (*θ_s_ -θ_t_*)/*θ_s_*, where *θ_s_* is the average simulation result and *θ_t_* is the theoretical expectation. The ± sign introduces a standard error. Statistically significant deviations from 0 are highlighted. ARGON_10_ results were obtained running ARGON with non-recombinant blocks of 10*μM*.

| | $n = N_e$ | singletons % | doubletons % |
| --- | --- | --- | --- |
| MS | 20,000 | **$-10.81 \pm 0.01$** | **$+4.87 \pm 0.02$** |
| ARGON | 20,000 | $+0.01 \pm 0.01$ | $-0.02 \pm 0.02$ |
| ARGON<sub>10</sub> | 20,000 | $+0.00 \pm 0.01$ | $-0.02 \pm 0.02$ |
| MS | 1,000 | **$-10.80 \pm 0.05$** | **$+4.80 \pm 0.09$** |
| ARGON | 1,000 | $+0.03 \pm 0.05$ | $-0.11 \pm 0.09$ |
| ARGON<sub>10</sub> | 1,000 | $+0.02 \pm 0.05$ | $+0.17 \pm 0.09$ |

### 3.2 Deviation of the coalescent from the DTWF

The coalescent becomes a poor approximation of the DTWF process when the sample size is not substantially smaller than the effective population size ([Wakeley and Takahashi, 2003, Bhaskar et al., 2014]). We verified that ARGON matches the theoretical prediction for the number of singletons and doubletons described in [Bhaskar et al., 2014] for the DTWF (see Table 1). We simulated populations of effective size *N_e_* = 1,000 and *N_e_* = 20,000 haploid individuals, and sampled all present-day individuals. While ARGON matches the prediction of [Bhaskar et al., 2014] in both exact and approximate mode, MS simulations substantially deviate from the DTWF model. Simulation using other coalescent approaches (e.g. COSI2, SCRM, MSprime), will similarly deviate in these scenarios.

### 3.3 Scalability to large sample size and whole-chromosome length

We tested the run time and memory usage of ARGON and two recently developed programs that enable simulating very large sample sizes and long chromosomes: COSI2 and SCRM (see Table 2). We find that for large parameter values SCRM generally performs worse than ARGON. COSI2 is generally faster, but scales poorly for memory usage as the size of the region and the sample size grows. Additional tests, including a pre-release version of MSprime, are detailed in the Supplementary Note.

We further tested the performance of approximate algorithms for the same set of simulation parameters (see Table 3). We compared ARGON with a minimum recombination block size of 10*μM* (AR_10_), SCRM with the “-l” flag set to 0 (SC_0_), and COSI2 with the “-u” flag set to 0 (CS_0_). We find that ARGON’s speed and memory usage is substantially improved, at the cost of slightly inflated correlation for neighboring markers (see Supplementary Note). For a constant population of size *N_e_* = 20,000, for instance, squared correlation (*r*^2^) of markers 0 to 50 Kb apart (using 1 cM/Mb) was increased by ~27% for AR_10_, but remained unchanged for markers at a larger distance. For CS_0_ and SC_0_, *r*^2^ was decreased by ~5% for markers at a larger distance. AR_10_ is faster than SC_0_, and approximately as fast as CS_0_, and uses less memory than both simulators for the large values of the test parameters. Comparison to other simulators and additional tests are detailed in the Supplementary Note.

**Table 2:** Comparison of simulation algorithms (AR=ARGON, SC=SCRM, CS=COSI2). Simulation parameters were the chromosome length (Mb), exponential expansion rate (measured as *ρ* = log(*A/C*)/*G* where *A* = ancestral size, *C* = current size, *G* = 500 = generation of expansion start), number of haploid samples (in thousands), and haploid ancestral population size (*A*). Recombination and mutation rates were set to 1 cM/Mb and 2 × 10^−8^ mutations per bp per generation, respectively. We compare simulation time (in hours) and memory usage (in Gb). All tests were run on a 2.27GHz Intel Xeon L5640, using a single core and up to 60 Gb of memory. † represents runs terminated due to insufficient memory (> 60Gb) or a memory error. Additional results are shown in the Supplementary Note.

| Mb | $10^2\rho$ | $n/10^3$ | A | Time | | | Memory | | |
| --- | --- | --- | --- | --- | --- | --- | --- | --- | --- |
|  |  |  |  | AR | SC | CS | AR | SC | CS |
| 100 | 0.0 | 20 | 20 | 0.48 | 1.42 | 0.12 | 10.1 | 19.6 | 17.3 |
| 300 | 0.0 | 20 | 20 | 2.56 | † | 0.59 | 15.2 | † | 41.4 |
| 500 | 0.0 | 20 | 20 | 5.35 | † | † | 19.8 | † | † |
| 100 | 0.0 | 1 | 10 | 0.09 | 0.02 | 0.01 | 6.5 | 0.5 | 0.2 |
| 100 | 0.0 | 10 | 10 | 0.12 | 0.27 | 0.02 | 8.5 | 4.9 | 4.4 |
| 100 | 0.0 | 100 | 10 | 1.01 | 6.20 | 52.33 | 12.6 | 48.9 | 6.8 |
| 100 | 0.0 | 200 | 10 | 3.05 | † | † | 19.4 | † | † |
| 100 | 0.0 | 300 | 10 | 10.19 | † | † | 24.6 | † | † |
| 100 | 0.347 | 20 | 10 | 0.39 | 1.37 | 0.07 | 10.5 | 19.6 | 9.0 |
| 100 | 0.576 | 20 | 10 | 0.94 | 2.37 | † | 13.0 | 39.1 | † |
| 100 | 1.151 | 20 | 10 | 3.97 | † | † | 25.0 | † | † |

**Table 3:** Comparison of approximate simulation algorithms (AR_10_=ARGON with minimum recombination block of size 10*μM*, SC_0_=SCRM with “-l” set to 0, CS_0_=COSI2 with “-u” set to 0). See Table 2 for a description of the simulation setup. † represents runs terminated due to insufficient memory (> 60Gb) or a memory error. ⋆ indicates that the program took longer than 100 hours to complete. Additional results are shown in the Supplementary Note.

| Mb | $10^2\rho$ | $n/10^3$ | A | Time | | | Memory | | |
| --- | --- | --- | --- | --- | --- | --- | --- | --- | --- |
|  |  |  |  | AR<sub>10</sub> | SC<sub>0</sub> | CS<sub>0</sub> | AR<sub>10</sub> | SC<sub>0</sub> | CS<sub>0</sub> |
| 100 | 0.0 | 20 | 20 | 0.12 | 1.23 | 0.15 | 8.2 | 19.6 | 17.3 |
| 300 | 0.0 | 20 | 20 | 0.46 | † | 0.89 | 9.7 | † | 41.4 |
| 500 | 0.0 | 20 | 20 | 0.79 | † | † | 11.0 | † | † |
| 100 | 0.0 | 1 | 10 | 0.02 | 0.02 | 0.01 | 3.3 | 0.5 | 0.3 |
| 100 | 0.0 | 10 | 10 | 0.04 | 0.28 | 0.03 | 7.9 | 4.9 | 4.4 |
| 100 | 0.0 | 100 | 10 | 0.34 | 6.36 | ★ | 8.0 | 48.9 | ★ |
| 100 | 0.0 | 200 | 10 | 0.66 | † | ★ | 8.4 | † | ★ |
| 100 | 0.0 | 300 | 10 | 1.59 | † | † | 17.6 | † | † |
| 100 | 0.347 | 20 | 10 | 0.09 | 2.51 | 0.07 | 8.2 | 39.1 | 8.9 |
| 100 | 0.576 | 20 | 10 | 0.15 | † | 0.13 | 8.6 | † | 9.4 |
| 100 | 1.151 | 20 | 10 | 0.22 | † | † | 12.0 | † | † |

## 4 Discussion

We described ARGON, a fast and scalable simulator for the discrete time Wright-Fisher process. Accurate simulation of large genealogical and sequence data will facilitate analysis of modern day genomic data sets, which have now reached hundreds of thousands of samples ([Sudlow et al., 2015]). ARGON has similar or superior time and memory performance compared to current coalescent simulators when large samples and long chromosomes are simulated, and enables accurately simulating the DTWF process when the sample size approaches the effective population size, where the coalescent approximation makes current scalable simulators become imprecise. Version 0.1 of ARGON supports arbitrary demographic history with migration, variable mutation/recombination rate and gene conversion, and outpus Newick trees and pairwise IBD segments for large sample sizes in both exact and approximate modes.

## Acknowledgements

We thank Alkes L. Price and John Wakeley for useful discussions and comments on an early draft.

## Supplementary Note

### 1 Approach

**Sampling the ARG**: ARGON proceeds backwards in time one generation at a time, occasionally sampling coalescent and recombination events subject to population structure and migration. Each individual is represented as a list of regions that are still being tracked at the current time (i.e. for which not all samples have found a common ancestor). Recombination events are sampled in genetic space from an exponential distribution, and rounded to the closest physiscal base pair position based on the desired recombination rate, which may be varying along the chromosome. A recombination event can be a crossover or a non-crossover event, based on user-specified rates. For each individual, a maximum of two parents are sampled, so that multiple recombination events result in alternating ancestry between two individuals from the previous generation. When two or more individuals choose the same parent, coalescence occurs if the individuals contain overlapping regions of genetic material. During coalescence, regions within individuals are merged and annotated with links to descendant ancestral recombination graph (ARG [Griffiths, 1981]) nodes.

**Sampling sequences from the ARG**: Once the ARG has been computed, mutation events are sampled on the graph edges according to a Poisson process. For each ARG edge affected by mutation, it is neccessary to calculate and store a list of samples that will be used to output genotypic information. Naïve calculation of these lists is intensive, and the storage of descendant lists for all nodes in the ARG results in prohibitive memory requirements. To overcome these difficulties, ARGON efficiently computes descendant lists using dynamic programming, and relies on a combination of top-down and bottom-up traversals of the ARG in order to always store the smallest possible set of descendant lists in memory while sampling mutations. During these operations, sparsity is exploited to further reduce memory consumption. Descendant lists are stored as either bit-set data structures, or lists of integers representing sample IDs, depending on their size. During the ARG traversals, whenever the size of a descendant list grows to exceed the expected memory consumption of the corresponding bit-set representation, the list is converted to the bit-set format. Similarly, ARGON offers an option (“-shrink” flag) to output the genotypes of the simulated individuals using a representation where the carriers of a derived allele are reported as a list of sample IDs, rather than the classical 0/1 representation of ancestral/derived alleles. Because a large number of mutations will have a set number of carriers, this representation saves computational speed and output disk space, and should be used to increase performance. Output formats also include the VCF format (default) and the hap/samples format.

These and other algorithmic details enable ARGON to be substantially more efficent than other DTWF implementations. In the GENOME simulator, for instance, individuals are represented as arrays of blocks of genetic material of a fixed size, and individuals from all populations are explicitly represented in memory using arrays. In ARGON, large regions are represented as intervals with arbitrary boundary values, and hash map data structures are extensively used to take advantage of sparsity, avoiding explicit representation of all individuals. As in the GENOME simulator, ARGON can run in approximate mode, so that recombinations breakpoints are rounded to blocks of a user-specified genetic length. This reduces the granularity of the recombination process, improving speed and memory usage, at the cost of slightly inflated correlation of markers at a short genomic distance. We tested this approximation using non-recombinant blocks of size 10*μM* and 50*μM* (1*μM* = 0.000001 Morgans).

### 2 Time and memory performance

We report detailed time and memory performance evaluation results for several simulators. Table 1 shows results for ARGON version 0.1.160101, SCRM version 1.6.1 [Staab et al., 2015], COSI version 2.0 [Shlyakhter et al., 2014], and MSprime version 0.1.6 [Kelleher et al., 2015]. ARGON was run using the “-shrink” flag and writing genotypes in the “seq” format, in which rare alleles are represented as lists of sample IDs. MSprime was run using the “-max-memory=60G” flag to enable generating large genealogies. Table 2 reports performance evaluation for several algorithms run in approximate mode: ARGON version 0.1.160101 with 10 and 50 microMorgans, FAST-SIMCOAL version 2.5.2.21 (implementing the SMC’ approximation), SCRM version 1.6.1 with the “-l” flag set to 0, and COSI version 2.0 with the “-u” flag set to 0. Note that the approximation flags for COSI2 and SCRM were set to the smallest allowed value, and that for both simulators slightly larger values may be used to find the appropriate trade-off between accuracy and speed. Table 3 reports a comparison of ARGON 0.1.160101 and GENOME version 0.2 [Liang et al., 2007], both run with a non-recombinant block size of 10 and 50 microMorgans. GENOME could not be run in exact mode due to excessive time and memoty requirements.

**Table 1:** Comparison of simulation algorithms (AR=ARGON, SC=SCRM, CS=COSI2, MS’=MSprime). Simulation parameters were the chromosome length (Mb), exponential expansion rate (measured as *ρ* = log(*A/C*)/*G* where *A* = ancestral size, *C* = current size, *G* = 500 = generation of expansion start), number of haploid samples (in thousands), and haploid ancestral population size (A). Recombination and mutation rates were set to 1 cM/Mb and 2 × 10^−8^ mutations per bp per generation, respectively. We compare simulation time (in hours) and memory usage (in Gb). All tests were run on a 2.27GHz Intel Xeon L5640, using a single core and up to 60 Gb of memory. † represents runs terminated due to insufficient memory (> 60Gb) or a memory error.

| Mb | $\rho$ | $n/10^3$ | A | Time | | | | Mem | | | |
| --- | --- | --- | --- | --- | --- | --- | --- | --- | --- | --- | --- |
|  |  |  |  | AR | SC | CS | MS' | AR | SC | CS | MS' |
| 100 | 0.0 | 20 | 20 | 0.48 | 1.42 | 0.12 | 0.05 | 10.1 | 19.6 | 17.3 | 2.3 |
| 300 | 0.0 | 20 | 20 | 2.56 | † | 0.59 | 0.27 | 15.2 | † | 41.4 | 6.8 |
| 500 | 0.0 | 20 | 20 | 5.35 | † | † | 0.75 | 19.8 | † | † | 11.3 |
| 700 | 0.0 | 20 | 20 | 10.16 | † | † | 1.45 | 24.7 | † | † | 15.7 |
| 900 | 0.0 | 20 | 20 | 21.63 | † | † | 2.47 | 24.8 | † | † | 20.2 |
| 100 | 0.0 | 1 | 10 | 0.09 | 0.02 | 0.01 | 0.01 | 6.5 | 0.5 | 0.2 | 0.1 |
| 100 | 0.0 | 10 | 10 | 0.12 | 0.27 | 0.02 | 0.01 | 8.5 | 4.9 | 4.4 | 0.6 |
| 100 | 0.0 | 100 | 10 | 1.01 | 6.20 | 52.33 | 0.10 | 12.6 | 48.9 | 6.8 | 5.8 |
| 100 | 0.0 | 200 | 10 | 3.05 | † | † | 0.28 | 19.4 | † | † | 12.2 |
| 100 | 0.0 | 300 | 10 | 10.19 | † | † | 0.38 | 24.6 | † | † | 18.7 |
| 100 | 0.00173 | 20 | 10 | 0.28 | 0.97 | 0.06 | 0.03 | 9.4 | 19.6 | 8.8 | 1.7 |
| 100 | 0.00347 | 20 | 10 | 0.39 | 1.37 | 0.07 | 0.05 | 10.5 | 19.6 | 9.0 | 2.4 |
| 100 | 0.00448 | 20 | 10 | 0.60 | 1.90 | 0.08 | 0.05 | 11.5 | 39.1 | 9.1 | 3.1 |
| 100 | 0.00520 | 20 | 10 | 0.78 | 2.10 | 0.44 | 0.06 | 12.3 | 39.1 | 56.6 | 3.6 |
| 100 | 0.00576 | 20 | 10 | 0.94 | 2.37 | † | 0.08 | 13.0 | 39.1 | † | 4.1 |
| 100 | 0.01151 | 20 | 10 | 3.97 | † | † | 0.59 | 25.0 | † | † | 14.4 |

**Table 2:** Comparison of approximate algorithms (AR_10_=ARGON with 10*μM* blocks, AR_50_=ARGON with 50*μM* blocks, FS=FASTSIMCOAL, SC_0_=SCRM with “-l” flag set to 0, CS_0_=COSI2 with “-u” flag set to 0), time and memory performance. Simulation parameters were the chromosome length (Mb), exponential expansion rate (measured as *ρ* = log(*A/C*)/*G* where *A* = ancestral size, *C* = current size, *G* = 500 = generation of expansion start), number of haploid samples (in thousands), and haploid ancestral population size (A). Recombination and mutation rates were set to 1 cM/Mb and 2 × 10^−8^ mutations per bp per generation, respectively. We compare simulation time (in hours) and memory usage (in Gb). All tests were run on a 2.27GHz Intel Xeon L5640, using a single core and up to 60 Gb of memory. † represents runs terminated due to insufficient memory (> 60Gb) or a memory error. (⋆) indicates that the program took longer than 100 hours to complete. * indicates that the result has been linearly extrapolated from previous runs with smaller parameter values.

| Mb | $\rho$ | $n/10^3$ | A | Time | | | | | Mem | | | | |
| --- | --- | --- | --- | --- | --- | --- | --- | --- | --- | --- | --- | --- | --- |
|  |  |  |  | AR<sub>10</sub> | AR<sub>50</sub> | FS | SC<sub>0</sub> | CS<sub>0</sub> | AR<sub>10</sub> | AR<sub>50</sub> | FS | SC<sub>0</sub> | CS<sub>0</sub> |
| 100 | 0.0 | 20 | 20 | 0.12 | 0.05 | 4.34 | 1.23 | 0.15 | 8.2 | 8.0 | 7.5 | 19.6 | 17.3 |
| 300 | 0.0 | 20 | 20 | 0.46 | 0.18 | 13.69* | † | 0.89 | 9.7 | 8.7 | 7.5* | † | 41.4 |
| 500 | 0.0 | 20 | 20 | 0.79 | 0.36 | 22.83* | † | † | 11.0 | 9.7 | 7.5* | † | † |
| 700 | 0.0 | 20 | 20 | 1.52 | 0.55 | 31.97* | † | † | 12.4 | 10.2 | 7.5* | † | † |
| 900 | 0.0 | 20 | 20 | 1.91 | 0.65 | 41.11* | † | † | 13.5 | 10.9 | 7.5* | † | † |
| 100 | 0.0 | 1 | 10 | 0.02 | 0.03 | 0.03 | 0.02 | 0.01 | 3.3 | 1.6 | 0.4 | 0.5 | 0.3 |
| 100 | 0.0 | 10 | 10 | 0.04 | 0.02 | 0.94 | 0.28 | 0.03 | 7.9 | 5.3 | 3.8 | 4.9 | 4.4 |
| 100 | 0.0 | 100 | 10 | 0.34 | 0.12 | 13.63 | 6.36 | (★) | 8.0 | 4.3 | 37.4 | 48.9 | (★) |
| 100 | 0.0 | 200 | 10 | 0.66 | 0.37 | † | † | (★) | 8.4 | 10.4 | † | † | (★) |
| 100 | 0.0 | 300 | 10 | 1.59 | 1.01 | † | † | † | 17.6 | 19.4 | † | † | † |
| 100 | 0.00173 | 20 | 10 | 0.06 | 0.04 | 2.90 | 1.31 | 0.10 | 8.1 | 6.7 | 7.5 | 19.6 | 8.8 |
| 100 | 0.00347 | 20 | 10 | 0.09 | 0.04 | 3.24 | 2.51 | 0.07 | 8.2 | 8.0 | 7.5 | 39.1 | 8.9 |
| 100 | 0.00448 | 20 | 10 | 0.17 | 0.09 | 4.68 | † | 0.09 | 8.7 | 8.1 | 7.5 | † | 9.2 |
| 100 | 0.00520 | 20 | 10 | 0.14 | 0.14 | 5.45 | † | 0.09 | 8.9 | 8.0 | 7.5 | † | 9.2 |
| 100 | 0.00576 | 20 | 10 | 0.15 | 0.10 | 5.51 | † | 0.13 | 8.6 | 8.0 | 7.6 | † | 9.4 |
| 100 | 0.01151 | 20 | 10 | 0.22 | 0.12 | 18.90 | † | † | 12.0 | 9.9 | 7.6 | † | † |

**Table 3:** Comparison of approximate algorithms (AR_10_=ARGON with 10*μM* blocks, AR_50_=ARGON with 50*μM* blocks, GEN_10_=GENOME with 10*μM* blocks, GEN_50_=GENOME with 50*μM* blocks), time and memory performance. Simulation parameters were the chromosome length (Mb), exponential expansion rate (measured as *ρ* = log(*A/C*)/*G* where *A* = ancestral size, *C* = current size, *G* = 500 = generation of expansion start), number of haploid samples (in thousands), and haploid ancestral population size (A). Recombination and mutation rates were set to 1 cM/Mb and 2 × 10^−8^, respectively. We compare simulation time (in hours) and memory usage (in Gb). All tests were run on a 2.27GHz Intel Xeon L5640, using a single core and up to 60 Gb of memory. † represents runs terminated due to insufficient memory (> 60Gb) or a memory error.

| Mb | $\rho$ | $n/10^3$ | A | Time | | | | Mem | | | |
| --- | --- | --- | --- | --- | --- | --- | --- | --- | --- | --- | --- |
|  |  |  |  | AR<sub>10</sub> | AR<sub>50</sub> | GEN<sub>10</sub> | GEN<sub>50</sub> | AR<sub>10</sub> | AR<sub>50</sub> | GEN<sub>10</sub> | GEN<sub>50</sub> |
| 100 | 0.0 | 20 | 20 | 0.12 | 0.05 | 0.50 | 0.44 | 8.2 | 8.0 | 18.0 | 5.9 |
| 300 | 0.0 | 20 | 20 | 0.46 | 0.18 | 1.49 | 1.30 | 9.7 | 8.7 | 54.5 | 17.8 |
| 500 | 0.0 | 20 | 20 | 0.79 | 0.36 | † | 2.23 | 11.0 | 9.7 | † | 30.1 |
| 700 | 0.0 | 20 | 20 | 1.52 | 0.55 | † | 3.01 | 12.4 | 10.2 | † | 41.2 |
| 900 | 0.0 | 20 | 20 | 1.91 | 0.65 | † | 3.97 | 13.5 | 10.9 | † | 53.9 |
| 100 | 0.0 | 1 | 10 | 0.02 | 0.03 | 0.02 | 0.01 | 3.3 | 1.6 | 0.9 | 0.2 |
| 100 | 0.0 | 10 | 10 | 0.04 | 0.02 | 0.18 | 0.10 | 7.9 | 5.3 | 8.8 | 1.8 |
| 100 | 0.0 | 100 | 10 | 0.34 | 0.12 | † | 1.94 | 8.0 | 4.3 | † | 28.3 |
| 100 | 0.0 | 200 | 10 | 0.66 | 0.37 | † | † | 8.4 | 10.4 | † | † |
| 100 | 0.0 | 300 | 10 | 1.59 | 1.01 | † | † | 17.6 | 19.4 | † | † |
| 100 | 0.00173 | 20 | 10 | 0.06 | 0.04 | 0.36 | 0.38 | 8.1 | 6.7 | 17.5 | 5.5 |
| 100 | 0.00347 | 20 | 10 | 0.09 | 0.04 | 0.55 | 0.46 | 8.2 | 8.0 | 17.7 | 6.0 |
| 100 | 0.00448 | 20 | 10 | 0.17 | 0.09 | 0.71 | 0.62 | 8.7 | 8.1 | 17.8 | 7.5 |
| 100 | 0.00520 | 20 | 10 | 0.14 | 0.14 | 0.79 | 0.66 | 8.9 | 8.0 | 17.9 | 8.0 |
| 100 | 0.00576 | 20 | 10 | 0.15 | 0.10 | 0.86 | 0.83 | 8.6 | 8.0 | 18.8 | 9.2 |
| 100 | 0.01151 | 20 | 10 | 0.22 | 0.12 | 2.39 | 2.09 | 12.0 | 9.9 | 29.7 | 20.2 |

### 3 Simulator accuracy

We compared ARGON, MS, COSI2, SCRM and MSprime to test accuracy for small sample sizes, when the coalescent process is a good approximation of the discrete time Wright-Fisher process. We tested a number of scenarios, for which we provide a description and corresponding MS command below. We did not test all scenarios for MSprime, because version 0.1.6 does not support some of the required features.

- Constant size: A population of constant size *N* = 20,000 haploid individuals; 50 haploid samples of 5 Mb, 10^−8^ recombinations per bp per generation, 2 × 10^−8^ mutations per bp per generation.

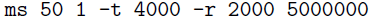
- Exponential expansion: A population of ancestral size *N* = 20, 000 haploid individuals, expanding from generation 500 to a present day size of 200,000 haploid individuals; 50 haploid samples of 5 Mb, 10^−8^ recombinations per bp per generation, 2 × 10^−8^ mutations per bp per generation.

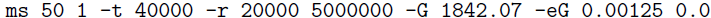
- Bottleneck: A population of ancestral size *N* = 20, 000 haploid individuals, shrinking from generation 500 to a present day size of 2, 000 haploid individuals; 50 haploid samples of 5 Mb, 10^−8^ recombinations per bp per generation, 2 × 10^−8^ mutations per bp per generation.

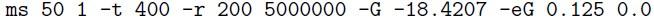
- Island model: 4 demes of constant size 5,000 haploid individuals, symmetric pairwise migration rate of 0.001 individuals per generation; 20 samples of 5 Mb from each deme, 10^−8^ recombinations per bp per generation, 2 × 10^−8^ mutations per bp per generation.

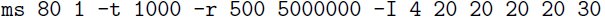
- Island model, high migration: 4 demes of constant size 5000 haploid individuals, symmetric pairwise migration rate of 0.01 individuals per generation; 20 samples of 5 Mb from each deme, 10^−8^ recombinations per bp per generation, 2 × 10^−8^ mutations per bp per generation.

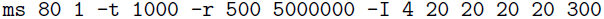
- Piecewise constant: A population of varying constant size. Size 100, 000 haploid individuals from generation 0 to 100; 5, 000 individuals from generation 100 to 200; 50, 000 haploid individuals from generation 200 to 300; and ancestral size 10, 000 haploid individuals from generation 300 on; 50 haploid samples of 5 Mb, 10^−8^ recombinations per bp per generation, 2 × 10^−8^ mutations per bp per generation.

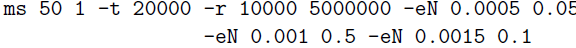

The flag “-c 1 300” was added when simulating gene conversion with same rate as recombination, and a mean tract length of 300 bp. The flag “-p 10” was added in all cases to increse the number of reported significant digits, and “-seed” was used to generate independent random draws. For the COSI2 simulator the infinite sites model was enabled adding the “infinite_sites yes” command in the configuration file. We tested the following summary statistics:

- *r*_50_ correlation of markers at distance between 0 and 50 Kb.
- *r*_00_ correlation of markers at distance between 50 and 100 Kb.
- *r*_150_ correlation of markers at distance between 100 and 150 Kb.
- 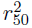 squared correlation of markers at distance between 0 and 50 Kb.
- 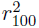 squared correlation of markers at distance between 50 and 100 Kb.
- 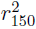 squared correlation of markers at distance between 100 and 150 Kb.
- *θ* heterozygosity computed as 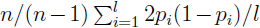 for *n* samples and *l* bases in the region, each with sample frequency *p_i_*.
- *S* number of polymorphic sites in the sample.
- *S*_<.1_ number of polymorphic sites with minor allele frequency below 10% in the sample.
- *S*_∈[.1,.4]_ number of polymorphic sites with minor allele frequency between 10% and 40% in the sample.
- *S*_>.4_ number of polymorphic sites with minor allele frequency above 40% in the sample.

*r* and *r*^2^ statistics were obtained using Plink 1.9 using the flags “-r” and “-r2”, for variants with MAF greater than 10%. The tables below report the value of these statistics for the considered pairs of simulators after 5, 000 independent draws. The column Δ% reports differences of each feature as 100 × (*mean*_1_ — *mean*_2_)/*mean*_1_. Z-scores corresponding to *p* < 0.001 are highlighted with an asterisk. Results are reported in tables 4 through 52.

We tested the accuracy of ARGON when a recombination map is used in input, and identical-by-descent (IBD) segments are requested in output. We simulated 400 haploid samples from populations of constant effective size ranging from 5, 000 to 30, 000 haploid individuals. We performed 10 independent simulations of a realistic ~ 249 Mb Human Chromosome 1, using the recombination map downloaded from https://mathgen.stats.ox.ac.uk/impute/1000GP_Phase3.html. To test the accuracy of the IBD provided in output, we estimated the size of the simulated populations using output segments of length greater than 2 cM. To estimate the effective population size, we used the maximum likelihood estimator derived in [Palamara, 2014]:

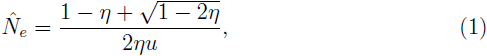

where 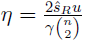 and *Ŝ_R_* represents the total number of segments longer than *u*

Morgans observed for all 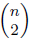 pairs. γ represents the total length (in Morgans) of the analyzed genomes. We observed good agreement between inferred and simulated population sizes (Figure 1), indicating that the distribution of IBD segments output by ARGON matches theoretical expectations. We plotted the density of recombination events, which we estimated using all edges of the IBD segments output across all simulations. As shown in figures 2 and 3, the density of observed recombination events follows the recombination map specified in input.

**Figure 1:**
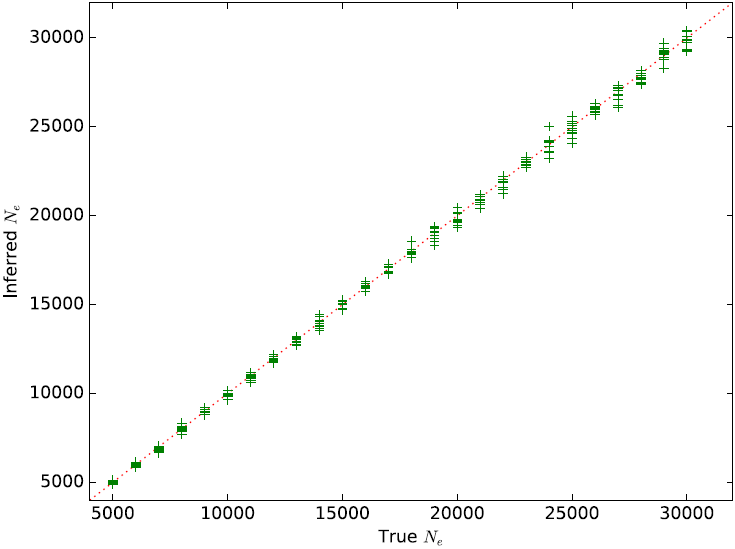
Comparison of simulated effective population size vs. the population size estimated using Eq. 1 and IBD segments of length at least 2 cM output by ARGON.

**Figure 2:**
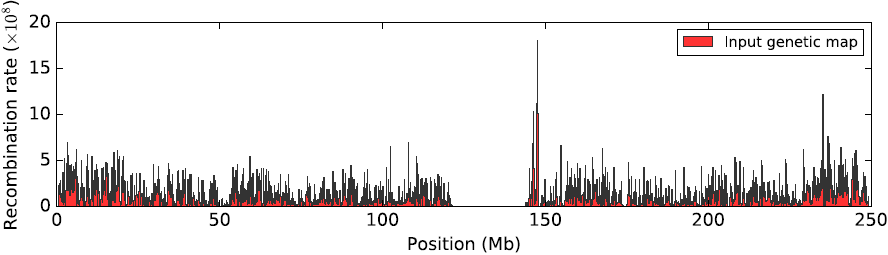
Genetic map.

**Figure 3:**
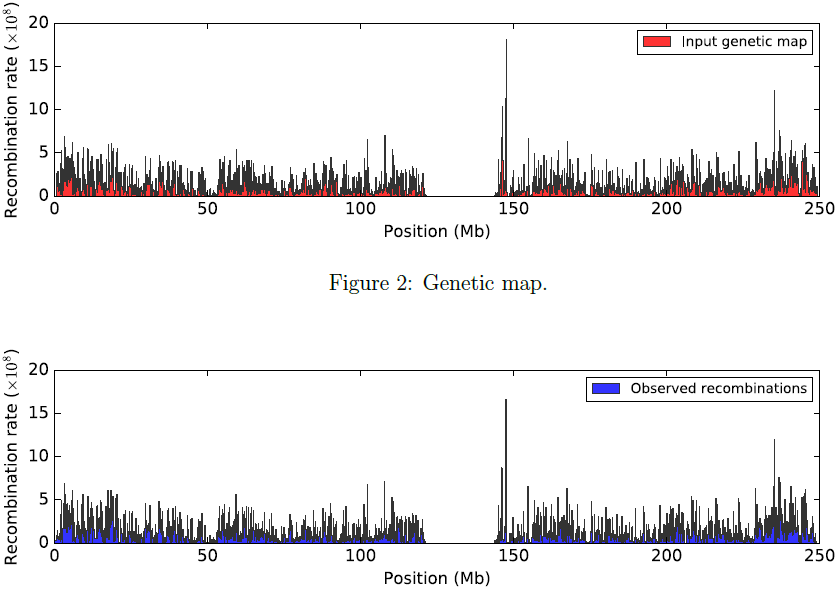
Density of IBD segment edges.

#### 3.1 ARGON and MS

**Table 4:** *s*_1_ = ARGON, *s*_2_ = MS; model: constant size

| feature | $s_1$ mean | $s_1$ s.e. | $s_2$ mean | $s_2$ s.e. | $\Delta\%$ | z-score |
| --- | --- | --- | --- | --- | --- | --- |
| $r_{50}$ | $9.2195 \times 10^{-2}$ | $3.5 \times 10^{-5}$ | $9.2157 \times 10^{-2}$ | $3.5 \times 10^{-5}$ | 0.0 | 0.77 |
| $r_{100}$ | $5.8528 \times 10^{-2}$ | $2.1 \times 10^{-5}$ | $5.8524 \times 10^{-2}$ | $2.1 \times 10^{-5}$ | 0.0 | 0.13 |
| $r_{150}$ | $5.2589 \times 10^{-2}$ | $1.9 \times 10^{-5}$ | $5.2587 \times 10^{-2}$ | $1.8 \times 10^{-5}$ | 0.0 | 0.08 |
| $r_{50}^2$ | $2.2082 \times 10^{-2}$ | $3.6 \times 10^{-5}$ | $2.2059 \times 10^{-2}$ | $3.6 \times 10^{-5}$ | 0.1 | 0.45 |
| $r_{100}^2$ | $3.1710 \times 10^{-3}$ | $1.8 \times 10^{-5}$ | $3.1556 \times 10^{-3}$ | $1.8 \times 10^{-5}$ | 0.5 | 0.60 |
| $r_{150}^2$ | $1.6683 \times 10^{-3}$ | $1.5 \times 10^{-5}$ | $1.6424 \times 10^{-3}$ | $1.6 \times 10^{-5}$ | 1.6 | 1.18 |
| $\theta$ | $7.9992 \times 10^{-4}$ | $3.4 \times 10^{-7}$ | $7.9966 \times 10^{-4}$ | $3.4 \times 10^{-7}$ | 0.0 | 0.54 |
| S | $1.7911 \times 10^4$ | 6.1 | $1.7916 \times 10^4$ | 6.1 | -0.0 | -0.58 |
| $S_{<.1}$ | $9.5553 \times 10^3$ | 4.4 | $9.5639 \times 10^3$ | 4.4 | -0.1 | -1.38 |
| $S_{\in[.1,.4]}$ | $6.5648 \times 10^3$ | 3.5 | $6.5642 \times 10^3$ | 3.4 | 0.0 | 0.12 |
| $S_{>.4}$ | $1.7905 \times 10^3$ | 1.8 | $1.7879 \times 10^3$ | 1.8 | 0.1 | 1.02 |

**Table 5:** *s*_1_ = ARGON, *s*_2_ = MS; model: exponential expansion

| feature | $s_1$ mean | $s_1$ s.e. | $s_2$ mean | $s_2$ s.e. | $\Delta\%$ | z-score |
| --- | --- | --- | --- | --- | --- | --- |
| $r_{50}$ | $8.1185 \times 10^{-2}$ | $2.4 \times 10^{-5}$ | $8.1150 \times 10^{-2}$ | $2.4 \times 10^{-5}$ | 0.0 | 1.03 |
| $r_{100}$ | $5.1489 \times 10^{-2}$ | $1.2 \times 10^{-5}$ | $5.1469 \times 10^{-2}$ | $1.2 \times 10^{-5}$ | 0.0 | 1.18 |
| $r_{150}$ | $4.6677 \times 10^{-2}$ | $9.5 \times 10^{-6}$ | $4.6659 \times 10^{-2}$ | $9.5 \times 10^{-6}$ | 0.0 | 1.34 |
| $r_{50}^2$ | $1.8069 \times 10^{-2}$ | $2.8 \times 10^{-5}$ | $1.8049 \times 10^{-2}$ | $2.9 \times 10^{-5}$ | 0.1 | 0.50 |
| $r_{100}^2$ | $2.0460 \times 10^{-3}$ | $1.3 \times 10^{-5}$ | $2.0228 \times 10^{-3}$ | $1.3 \times 10^{-5}$ | 1.1 | 1.26 |
| $r_{150}^2$ | $8.6980 \times 10^{-4}$ | $1.1 \times 10^{-5}$ | $8.5858 \times 10^{-4}$ | $1.1 \times 10^{-5}$ | 1.3 | 0.72 |
| $\theta$ | $8.1197 \times 10^{-4}$ | $3.4 \times 10^{-7}$ | $8.1180 \times 10^{-4}$ | $3.4 \times 10^{-7}$ | 0.0 | 0.35 |
| S | $1.9363 \times 10^4$ | 5.8 | $1.9370 \times 10^4$ | 5.8 | -0.0 | -0.85 |
| $S_{<.1}$ | $1.1007 \times 10^4$ | 4.0 | $1.1017 \times 10^4$ | 4.0 | -0.1 | -1.77 |
| $S_{\in[.1,.4]}$ | $6.5682 \times 10^3$ | 3.4 | $6.5651 \times 10^3$ | 3.4 | 0.0 | 0.64 |
| $S_{>.4}$ | $1.7880 \times 10^3$ | 1.8 | $1.7877 \times 10^3$ | 1.8 | 0.0 | 0.12 |

**Table 6:** *s*_1_ = ARGON, *s*_2_ = MS; model: bottleneck

| feature | $s_1$ mean | $s_1$ s.e. | $s_2$ mean | $s_2$ s.e. | $\Delta\%$ | z-score |
| --- | --- | --- | --- | --- | --- | --- |
| $r_{50}$ | $1.4323 \times 10^{-1}$ | $1.0 \times 10^{-4}$ | $1.4335 \times 10^{-1}$ | $1.0 \times 10^{-4}$ | -0.1 | -0.85 |
| $r_{100}$ | $9.5918 \times 10^{-2}$ | $7.7 \times 10^{-5}$ | $9.5984 \times 10^{-2}$ | $7.7 \times 10^{-5}$ | -0.1 | -0.61 |
| $r_{150}$ | $8.5765 \times 10^{-2}$ | $7.2 \times 10^{-5}$ | $8.5805 \times 10^{-2}$ | $7.1 \times 10^{-5}$ | -0.0 | -0.40 |
| $r_{50}^2$ | $4.3686 \times 10^{-2}$ | $1.0 \times 10^{-4}$ | $4.3746 \times 10^{-2}$ | $9.9 \times 10^{-5}$ | -0.1 | -0.43 |
| $r_{100}^2$ | $1.1734 \times 10^{-2}$ | $6.4 \times 10^{-5}$ | $1.1788 \times 10^{-2}$ | $6.7 \times 10^{-5}$ | -0.5 | -0.58 |
| $r_{150}^2$ | $8.3447 \times 10^{-3}$ | $5.6 \times 10^{-5}$ | $8.3878 \times 10^{-3}$ | $5.7 \times 10^{-5}$ | -0.5 | -0.54 |
| $\theta$ | $7.4407 \times 10^{-4}$ | $3.7 \times 10^{-7}$ | $7.4371 \times 10^{-4}$ | $3.6 \times 10^{-7}$ | 0.0 | 0.70 |
| S | $1.3966 \times 10^4$ | 6.8 | $1.3956 \times 10^4$ | 6.9 | 0.1 | 1.03 |
| $S_{<.1}$ | $5.7458 \times 10^3$ | 5.1 | $5.7377 \times 10^3$ | 5.2 | 0.1 | 1.11 |
| $S_{\in[.1,.4]}$ | $6.4321 \times 10^3$ | 4.0 | $6.4319 \times 10^3$ | 3.9 | 0.0 | 0.04 |
| $S_{>.4}$ | $1.7883 \times 10^3$ | 1.9 | $1.7860 \times 10^3$ | 1.8 | 0.1 | 0.88 |

**Table 7:** *s*_1_ = ARGON, *s*_2_ = MS; model: island model

| feature | $s_1$ mean | $s_1$ s.e. | $s_2$ mean | $s_2$ s.e. | $\Delta\%$ | z-score |
| --- | --- | --- | --- | --- | --- | --- |
| $r_{50}$ | $7.1086 \times 10^{-2}$ | $2.7 \times 10^{-5}$ | $7.1124 \times 10^{-2}$ | $2.8 \times 10^{-5}$ | -0.1 | -0.98 |
| $r_{100}$ | $4.1662 \times 10^{-2}$ | $1.5 \times 10^{-5}$ | $4.1675 \times 10^{-2}$ | $1.6 \times 10^{-5}$ | -0.0 | -0.59 |
| $r_{150}$ | $3.6234 \times 10^{-2}$ | $1.3 \times 10^{-5}$ | $3.6240 \times 10^{-2}$ | $1.3 \times 10^{-5}$ | -0.0 | -0.33 |
| $r_{50}^2$ | $1.8451 \times 10^{-2}$ | $2.8 \times 10^{-5}$ | $1.8496 \times 10^{-2}$ | $2.9 \times 10^{-5}$ | -0.2 | -1.12 |
| $r_{100}^2$ | $2.7551 \times 10^{-3}$ | $1.3 \times 10^{-5}$ | $2.7469 \times 10^{-3}$ | $1.3 \times 10^{-5}$ | 0.3 | 0.45 |
| $r_{150}^2$ | $1.4684 \times 10^{-3}$ | $1.1 \times 10^{-5}$ | $1.4771 \times 10^{-3}$ | $1.1 \times 10^{-5}$ | -0.6 | -0.56 |
| $\theta$ | $8.1542 \times 10^{-4}$ | $3.5 \times 10^{-7}$ | $8.1444 \times 10^{-4}$ | $3.4 \times 10^{-7}$ | 0.1 | 2.01 |
| S | $2.0174 \times 10^4$ | 6.5 | $2.0163 \times 10^4$ | 6.4 | 0.1 | 1.21 |
| $S_{<.1}$ | $1.1493 \times 10^4$ | 4.6 | $1.1493 \times 10^4$ | 4.7 | 0.0 | 0.00 |
| $S_{\in[.1,.4]}$ | $6.9197 \times 10^3$ | 3.7 | $6.9126 \times 10^3$ | 3.6 | 0.1 | 1.38 |
| $S_{>.4}$ | $1.7615 \times 10^3$ | 1.7 | $1.7576 \times 10^3$ | 1.8 | 0.2 | 1.58 |

**Table 8:** *s*_1_ = ARGON, *s*_2_ = MS; island model, high migration

| feature | $s_1$ mean | $s_1$ s.e. | $s_2$ mean | $s_2$ s.e. | $\Delta\%$ | z-score |
| --- | --- | --- | --- | --- | --- | --- |
| $r_{50}$ | $7.1134 \times 10^{-2}$ | $2.8 \times 10^{-5}$ | $7.1115 \times 10^{-2}$ | $2.8 \times 10^{-5}$ | 0.0 | 0.48 |
| $r_{100}$ | $4.1621 \times 10^{-2}$ | $1.5 \times 10^{-5}$ | $4.1623 \times 10^{-2}$ | $1.5 \times 10^{-5}$ | -0.0 | -0.09 |
| $r_{150}$ | $3.6151 \times 10^{-2}$ | $1.3 \times 10^{-5}$ | $3.6161 \times 10^{-2}$ | $1.3 \times 10^{-5}$ | -0.0 | -0.54 |
| $r_{50}^2$ | $1.8363 \times 10^{-2}$ | $2.9 \times 10^{-5}$ | $1.8346 \times 10^{-2}$ | $2.8 \times 10^{-5}$ | 0.1 | 0.42 |
| $r_{100}^2$ | $2.6744 \times 10^{-3}$ | $1.3 \times 10^{-5}$ | $2.6739 \times 10^{-3}$ | $1.3 \times 10^{-5}$ | 0.0 | 0.03 |
| $r_{150}^2$ | $1.4068 \times 10^{-3}$ | $1.1 \times 10^{-5}$ | $1.4173 \times 10^{-3}$ | $1.1 \times 10^{-5}$ | -0.7 | -0.67 |
| $\theta$ | $8.0130 \times 10^{-4}$ | $3.4 \times 10^{-7}$ | $8.0187 \times 10^{-4}$ | $3.4 \times 10^{-7}$ | -0.1 | -1.19 |
| S | $1.9844 \times 10^4$ | 6.3 | $1.9848 \times 10^4$ | 6.1 | -0.0 | -0.46 |
| $S_{<.1}$ | $1.1313 \times 10^4$ | 4.7 | $1.1310 \times 10^4$ | 4.6 | 0.0 | 0.46 |
| $S_{\in[.1,.4]}$ | $6.8029 \times 10^3$ | 3.5 | $6.8066 \times 10^3$ | 3.5 | -0.1 | -0.75 |
| $S_{>.4}$ | $1.7275 \times 10^3$ | 1.8 | $1.7310 \times 10^3$ | 1.7 | -0.2 | -1.41 |

**Table 9:**
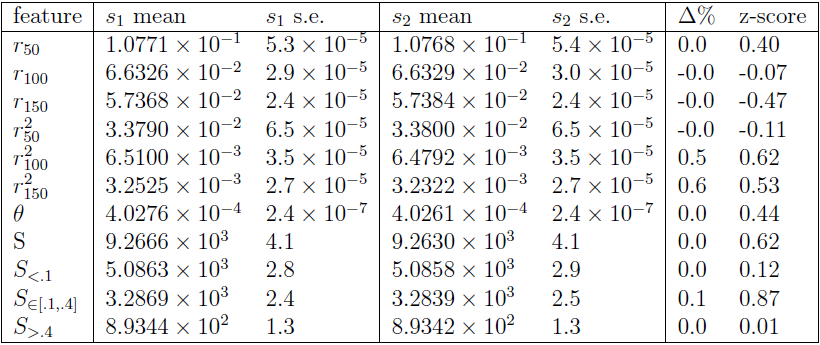
*s*_1_ = ARGON, *s*_2_ = MS; model: piecewise model

#### 3.2 ARGON and COSI2

**Table 10:**
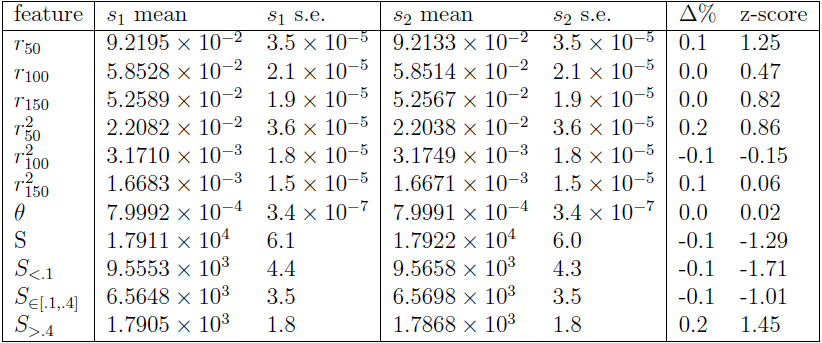
*s*_1_ = ARGON, *s*_2_ = COSI2; model: constant size

| feature | $s_1$ mean | $s_1$ s.e. | $s_2$ mean | $s_2$ s.e. | $\Delta\%$ | z-score |
| --- | --- | --- | --- | --- | --- | --- |
| $r_{50}$ | $9.2195 \times 10^{-2}$ | $3.5 \times 10^{-5}$ | $9.2133 \times 10^{-2}$ | $3.5 \times 10^{-5}$ | 0.1 | 1.25 |
| $r_{100}$ | $5.8528 \times 10^{-2}$ | $2.1 \times 10^{-5}$ | $5.8514 \times 10^{-2}$ | $2.1 \times 10^{-5}$ | 0.0 | 0.47 |
| $r_{150}$ | $5.2589 \times 10^{-2}$ | $1.9 \times 10^{-5}$ | $5.2567 \times 10^{-2}$ | $1.9 \times 10^{-5}$ | 0.0 | 0.82 |
| $r_{50}^2$ | $2.2082 \times 10^{-2}$ | $3.6 \times 10^{-5}$ | $2.2038 \times 10^{-2}$ | $3.6 \times 10^{-5}$ | 0.2 | 0.86 |
| $r_{100}^2$ | $3.1710 \times 10^{-3}$ | $1.8 \times 10^{-5}$ | $3.1749 \times 10^{-3}$ | $1.8 \times 10^{-5}$ | -0.1 | -0.15 |
| $r_{150}^2$ | $1.6683 \times 10^{-3}$ | $1.5 \times 10^{-5}$ | $1.6671 \times 10^{-3}$ | $1.5 \times 10^{-5}$ | 0.1 | 0.06 |
| $\theta$ | $7.9992 \times 10^{-4}$ | $3.4 \times 10^{-7}$ | $7.9991 \times 10^{-4}$ | $3.4 \times 10^{-7}$ | 0.0 | 0.02 |
| S | $1.7911 \times 10^4$ | 6.1 | $1.7922 \times 10^4$ | 6.0 | -0.1 | -1.29 |
| $S_{<.1}$ | $9.5553 \times 10^3$ | 4.4 | $9.5658 \times 10^3$ | 4.3 | -0.1 | -1.71 |
| $S_{\in[.1,.4]}$ | $6.5648 \times 10^3$ | 3.5 | $6.5698 \times 10^3$ | 3.5 | -0.1 | -1.01 |
| $S_{>.4}$ | $1.7905 \times 10^3$ | 1.8 | $1.7868 \times 10^3$ | 1.8 | 0.2 | 1.45 |

**Table 11:**
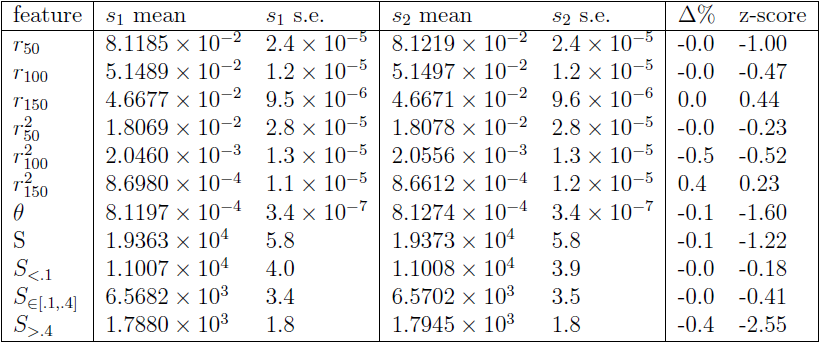
*s*_1_ = ARGON, *s*_2_ = COSI2; model: exponential expansion

| feature | $s_1$ mean | $s_1$ s.e. | $s_2$ mean | $s_2$ s.e. | $\Delta\%$ | z-score |
| --- | --- | --- | --- | --- | --- | --- |
| $r_{50}$ | $8.1185 \times 10^{-2}$ | $2.4 \times 10^{-5}$ | $8.1219 \times 10^{-2}$ | $2.4 \times 10^{-5}$ | -0.0 | -1.00 |
| $r_{100}$ | $5.1489 \times 10^{-2}$ | $1.2 \times 10^{-5}$ | $5.1497 \times 10^{-2}$ | $1.2 \times 10^{-5}$ | -0.0 | -0.47 |
| $r_{150}$ | $4.6677 \times 10^{-2}$ | $9.5 \times 10^{-6}$ | $4.6671 \times 10^{-2}$ | $9.6 \times 10^{-6}$ | 0.0 | 0.44 |
| $r_{50}^2$ | $1.8069 \times 10^{-2}$ | $2.8 \times 10^{-5}$ | $1.8078 \times 10^{-2}$ | $2.8 \times 10^{-5}$ | -0.0 | -0.23 |
| $r_{100}^2$ | $2.0460 \times 10^{-3}$ | $1.3 \times 10^{-5}$ | $2.0556 \times 10^{-3}$ | $1.3 \times 10^{-5}$ | -0.5 | -0.52 |
| $r_{150}^2$ | $8.6980 \times 10^{-4}$ | $1.1 \times 10^{-5}$ | $8.6612 \times 10^{-4}$ | $1.2 \times 10^{-5}$ | 0.4 | 0.23 |
| $\theta$ | $8.1197 \times 10^{-4}$ | $3.4 \times 10^{-7}$ | $8.1274 \times 10^{-4}$ | $3.4 \times 10^{-7}$ | -0.1 | -1.60 |
| S | $1.9363 \times 10^4$ | 5.8 | $1.9373 \times 10^4$ | 5.8 | -0.1 | -1.22 |
| $S_{<.1}$ | $1.1007 \times 10^4$ | 4.0 | $1.1008 \times 10^4$ | 3.9 | -0.0 | -0.18 |
| $S_{\in[.1,.4]}$ | $6.5682 \times 10^3$ | 3.4 | $6.5702 \times 10^3$ | 3.5 | -0.0 | -0.41 |
| $S_{>.4}$ | $1.7880 \times 10^3$ | 1.8 | $1.7945 \times 10^3$ | 1.8 | -0.4 | -2.55 |

**Table 12:** *s*_1_ = ARGON, *s*_2_ = COSI2; model: bottleneck

| feature | $s_1$ mean | $s_1$ s.e. | $s_2$ mean | $s_2$ s.e. | $\Delta\%$ | z-score |
| --- | --- | --- | --- | --- | --- | --- |
| $r_{50}$ | $1.4323 \times 10^{-1}$ | $1.0 \times 10^{-4}$ | $1.4333 \times 10^{-1}$ | $1.0 \times 10^{-4}$ | -0.1 | -0.71 |
| $r_{100}$ | $9.5918 \times 10^{-2}$ | $7.7 \times 10^{-5}$ | $9.5974 \times 10^{-2}$ | $8.0 \times 10^{-5}$ | -0.1 | -0.50 |
| $r_{150}$ | $8.5765 \times 10^{-2}$ | $7.2 \times 10^{-5}$ | $8.5847 \times 10^{-2}$ | $7.4 \times 10^{-5}$ | -0.1 | -0.79 |
| $r_{50}^2$ | $4.3686 \times 10^{-2}$ | $1.0 \times 10^{-4}$ | $4.3569 \times 10^{-2}$ | $9.8 \times 10^{-5}$ | 0.3 | 0.84 |
| $r_{100}^2$ | $1.1734 \times 10^{-2}$ | $6.4 \times 10^{-5}$ | $1.1653 \times 10^{-2}$ | $6.5 \times 10^{-5}$ | 0.7 | 0.89 |
| $r_{150}^2$ | $8.3447 \times 10^{-3}$ | $5.6 \times 10^{-5}$ | $8.2813 \times 10^{-3}$ | $5.6 \times 10^{-5}$ | 0.8 | 0.80 |
| $\theta$ | $7.4407 \times 10^{-4}$ | $3.7 \times 10^{-7}$ | $7.4432 \times 10^{-4}$ | $3.6 \times 10^{-7}$ | -0.0 | -0.48 |
| S | $1.3966 \times 10^4$ | 6.8 | $1.3962 \times 10^4$ | 6.8 | 0.0 | 0.42 |
| $S_{<.1}$ | $5.7458 \times 10^3$ | 5.1 | $5.7341 \times 10^3$ | 5.2 | 0.2 | 1.61 |
| $S_{\in[.1,.4]}$ | $6.4321 \times 10^3$ | 4.0 | $6.4396 \times 10^3$ | 4.0 | -0.1 | -1.33 |
| $S_{>.4}$ | $1.7883 \times 10^3$ | 1.9 | $1.7883 \times 10^3$ | 1.9 | 0.0 | 0.00 |

**Table 13:** *s*_1_ = ARGON, *s*_2_ = COSI2; model: island model

| feature | $s_1$ mean | $s_1$ s.e. | $s_2$ mean | $s_2$ s.e. | $\Delta\%$ | z-score |
| --- | --- | --- | --- | --- | --- | --- |
| $r_{50}$ | $7.1086 \times 10^{-2}$ | $2.7 \times 10^{-5}$ | $7.1052 \times 10^{-2}$ | $2.8 \times 10^{-5}$ | 0.0 | 0.87 |
| $r_{100}$ | $4.1662 \times 10^{-2}$ | $1.5 \times 10^{-5}$ | $4.1665 \times 10^{-2}$ | $1.5 \times 10^{-5}$ | -0.0 | -0.14 |
| $r_{150}$ | $3.6234 \times 10^{-2}$ | $1.3 \times 10^{-5}$ | $3.6246 \times 10^{-2}$ | $1.3 \times 10^{-5}$ | -0.0 | -0.65 |
| $r_{50}^2$ | $1.8451 \times 10^{-2}$ | $2.8 \times 10^{-5}$ | $1.8388 \times 10^{-2}$ | $2.9 \times 10^{-5}$ | 0.3 | 1.56 |
| $r_{100}^2$ | $2.7551 \times 10^{-3}$ | $1.3 \times 10^{-5}$ | $2.7295 \times 10^{-3}$ | $1.4 \times 10^{-5}$ | 0.9 | 1.34 |
| $r_{150}^2$ | $1.4684 \times 10^{-3}$ | $1.1 \times 10^{-5}$ | $1.4673 \times 10^{-3}$ | $1.1 \times 10^{-5}$ | 0.1 | 0.07 |
| $\theta$ | $8.1542 \times 10^{-4}$ | $3.5 \times 10^{-7}$ | $8.1551 \times 10^{-4}$ | $3.4 \times 10^{-7}$ | -0.0 | -0.18 |
| S | $2.0174 \times 10^4$ | 6.5 | $2.0183 \times 10^4$ | 6.3 | -0.0 | -0.99 |
| $S_{<.1}$ | $1.1493 \times 10^4$ | 4.6 | $1.1500 \times 10^4$ | 4.6 | -0.1 | -1.08 |
| $S_{\in[.1,.4]}$ | $6.9197 \times 10^3$ | 3.7 | $6.9251 \times 10^3$ | 3.6 | -0.1 | -1.05 |
| $S_{>.4}$ | $1.7615 \times 10^3$ | 1.7 | $1.7579 \times 10^3$ | 1.8 | 0.2 | 1.45 |

**Table 14:** *s*_1_ = ARGON, *s*_2_ = COSI2; model: island model, high migration

| feature | $s_1$ mean | $s_1$ s.e. | $s_2$ mean | $s_2$ s.e. | $\Delta\%$ | z-score |
| --- | --- | --- | --- | --- | --- | --- |
| $r_{50}$ | $7.1134 \times 10^{-2}$ | $2.8 \times 10^{-5}$ | $7.1094 \times 10^{-2}$ | $2.8 \times 10^{-5}$ | 0.1 | 1.01 |
| $r_{100}$ | $4.1621 \times 10^{-2}$ | $1.5 \times 10^{-5}$ | $4.1603 \times 10^{-2}$ | $1.5 \times 10^{-5}$ | 0.0 | 0.85 |
| $r_{150}$ | $3.6151 \times 10^{-2}$ | $1.3 \times 10^{-5}$ | $3.6156 \times 10^{-2}$ | $1.3 \times 10^{-5}$ | -0.0 | -0.27 |
| $r_{50}^2$ | $1.8363 \times 10^{-2}$ | $2.9 \times 10^{-5}$ | $1.8367 \times 10^{-2}$ | $2.8 \times 10^{-5}$ | -0.0 | -0.10 |
| $r_{100}^2$ | $2.6744 \times 10^{-3}$ | $1.3 \times 10^{-5}$ | $2.6820 \times 10^{-3}$ | $1.3 \times 10^{-5}$ | -0.3 | -0.41 |
| $r_{150}^2$ | $1.4068 \times 10^{-3}$ | $1.1 \times 10^{-5}$ | $1.4287 \times 10^{-3}$ | $1.1 \times 10^{-5}$ | -1.5 | -1.41 |
| $\theta$ | $8.0130 \times 10^{-4}$ | $3.4 \times 10^{-7}$ | $8.0114 \times 10^{-4}$ | $3.4 \times 10^{-7}$ | 0.0 | 0.33 |
| S | $1.9844 \times 10^4$ | 6.3 | $1.9842 \times 10^4$ | 6.3 | 0.0 | 0.22 |
| $S_{<.1}$ | $1.1313 \times 10^4$ | 4.7 | $1.1316 \times 10^4$ | 4.6 | -0.0 | -0.46 |
| $S_{\in[.1,.4]}$ | $6.8029 \times 10^3$ | 3.5 | $6.7946 \times 10^3$ | 3.6 | 0.1 | 1.65 |
| $S_{>.4}$ | $1.7275 \times 10^3$ | 1.8 | $1.7314 \times 10^3$ | 1.8 | -0.2 | -1.53 |

**Table 15:** *s*_1_ = ARGON, *s*_2_ = COSI2; model: piecewise model

| feature | $s_1$ mean | $s_1$ s.e. | $s_2$ mean | $s_2$ s.e. | $\Delta\%$ | z-score |
| --- | --- | --- | --- | --- | --- | --- |
| $r_{50}$ | $1.0771 \times 10^{-1}$ | $5.3 \times 10^{-5}$ | $1.0765 \times 10^{-1}$ | $5.4 \times 10^{-5}$ | 0.1 | 0.79 |
| $r_{100}$ | $6.6326 \times 10^{-2}$ | $2.9 \times 10^{-5}$ | $6.6282 \times 10^{-2}$ | $3.0 \times 10^{-5}$ | 0.1 | 1.05 |
| $r_{150}$ | $5.7368 \times 10^{-2}$ | $2.4 \times 10^{-5}$ | $5.7383 \times 10^{-2}$ | $2.4 \times 10^{-5}$ | -0.0 | -0.44 |
| $r_{50}^2$ | $3.3790 \times 10^{-2}$ | $6.5 \times 10^{-5}$ | $3.3773 \times 10^{-2}$ | $6.5 \times 10^{-5}$ | 0.1 | 0.18 |
| $r_{100}^2$ | $6.5100 \times 10^{-3}$ | $3.5 \times 10^{-5}$ | $6.4292 \times 10^{-3}$ | $3.4 \times 10^{-5}$ | 1.3 | 1.66 |
| $r_{150}^2$ | $3.2525 \times 10^{-3}$ | $2.7 \times 10^{-5}$ | $3.2436 \times 10^{-3}$ | $2.7 \times 10^{-5}$ | 0.3 | 0.23 |
| $\theta$ | $4.0276 \times 10^{-4}$ | $2.4 \times 10^{-7}$ | $4.0271 \times 10^{-4}$ | $2.5 \times 10^{-7}$ | 0.0 | 0.14 |
| S | $9.2666 \times 10^3$ | 4.1 | $9.2695 \times 10^3$ | 4.2 | -0.0 | -0.49 |
| $S_{<.1}$ | $5.0863 \times 10^3$ | 2.8 | $5.0921 \times 10^3$ | 2.9 | -0.1 | -1.44 |
| $S_{\in[.1,.4]}$ | $3.2869 \times 10^3$ | 2.4 | $3.2841 \times 10^3$ | 2.5 | 0.1 | 0.81 |
| $S_{>.4}$ | $8.9344 \times 10^2$ | 1.3 | $8.9321 \times 10^2$ | 1.3 | 0.0 | 0.13 |

#### 3.3 MS and COSI2

**Table 16:**
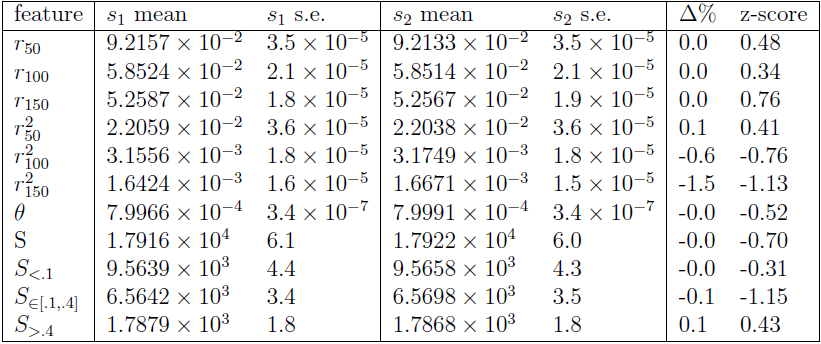
*s*_1_ = MS, *s*_2_ = COSI2; model: constant size

| feature | $s_1$ mean | $s_1$ s.e. | $s_2$ mean | $s_2$ s.e. | $\Delta\%$ | z-score |
| --- | --- | --- | --- | --- | --- | --- |
| $r_{50}$ | $9.2157 \times 10^{-2}$ | $3.5 \times 10^{-5}$ | $9.2133 \times 10^{-2}$ | $3.5 \times 10^{-5}$ | 0.0 | 0.48 |
| $r_{100}$ | $5.8524 \times 10^{-2}$ | $2.1 \times 10^{-5}$ | $5.8514 \times 10^{-2}$ | $2.1 \times 10^{-5}$ | 0.0 | 0.34 |
| $r_{150}$ | $5.2587 \times 10^{-2}$ | $1.8 \times 10^{-5}$ | $5.2567 \times 10^{-2}$ | $1.9 \times 10^{-5}$ | 0.0 | 0.76 |
| $r_{50}^2$ | $2.2059 \times 10^{-2}$ | $3.6 \times 10^{-5}$ | $2.2038 \times 10^{-2}$ | $3.6 \times 10^{-5}$ | 0.1 | 0.41 |
| $r_{100}^2$ | $3.1556 \times 10^{-3}$ | $1.8 \times 10^{-5}$ | $3.1749 \times 10^{-3}$ | $1.8 \times 10^{-5}$ | -0.6 | -0.76 |
| $r_{150}^2$ | $1.6424 \times 10^{-3}$ | $1.6 \times 10^{-5}$ | $1.6671 \times 10^{-3}$ | $1.5 \times 10^{-5}$ | -1.5 | -1.13 |
| $\theta$ | $7.9966 \times 10^{-4}$ | $3.4 \times 10^{-7}$ | $7.9991 \times 10^{-4}$ | $3.4 \times 10^{-7}$ | -0.0 | -0.52 |
| S | $1.7916 \times 10^4$ | 6.1 | $1.7922 \times 10^4$ | 6.0 | -0.0 | -0.70 |
| $S_{<.1}$ | $9.5639 \times 10^3$ | 4.4 | $9.5658 \times 10^3$ | 4.3 | -0.0 | -0.31 |
| $S_{\in[.1,.4]}$ | $6.5642 \times 10^3$ | 3.4 | $6.5698 \times 10^3$ | 3.5 | -0.1 | -1.15 |
| $S_{>.4}$ | $1.7879 \times 10^3$ | 1.8 | $1.7868 \times 10^3$ | 1.8 | 0.1 | 0.43 |

**Table 17:** *s*_1_ = MS, *s*_2_ = COSI2; model: exponential expansion

| feature | $s_1$ mean | $s_1$ s.e. | $s_2$ mean | $s_2$ s.e. | $\Delta\%$ | z-score |
| --- | --- | --- | --- | --- | --- | --- |
| $r_{50}$ | $8.1150 \times 10^{-2}$ | $2.4 \times 10^{-5}$ | $8.1219 \times 10^{-2}$ | $2.4 \times 10^{-5}$ | -0.1 | -2.03 |
| $r_{100}$ | $5.1469 \times 10^{-2}$ | $1.2 \times 10^{-5}$ | $5.1497 \times 10^{-2}$ | $1.2 \times 10^{-5}$ | -0.1 | -1.65 |
| $r_{150}$ | $4.6659 \times 10^{-2}$ | $9.5 \times 10^{-6}$ | $4.6671 \times 10^{-2}$ | $9.6 \times 10^{-6}$ | -0.0 | -0.89 |
| $r_{50}^2$ | $1.8049 \times 10^{-2}$ | $2.9 \times 10^{-5}$ | $1.8078 \times 10^{-2}$ | $2.8 \times 10^{-5}$ | -0.2 | -0.72 |
| $r_{100}^2$ | $2.0228 \times 10^{-3}$ | $1.3 \times 10^{-5}$ | $2.0556 \times 10^{-3}$ | $1.3 \times 10^{-5}$ | -1.6 | -1.78 |
| $r_{150}^2$ | $8.5858 \times 10^{-4}$ | $1.1 \times 10^{-5}$ | $8.6612 \times 10^{-4}$ | $1.2 \times 10^{-5}$ | -0.9 | -0.46 |
| $\theta$ | $8.1180 \times 10^{-4}$ | $3.4 \times 10^{-7}$ | $8.1274 \times 10^{-4}$ | $3.4 \times 10^{-7}$ | -0.1 | -1.95 |
| S | $1.9370 \times 10^4$ | 5.8 | $1.9373 \times 10^4$ | 5.8 | -0.0 | -0.37 |
| $S_{<.1}$ | $1.1017 \times 10^4$ | 4.0 | $1.1008 \times 10^4$ | 3.9 | 0.1 | 1.61 |
| $S_{\in[.1,.4]}$ | $6.5651 \times 10^3$ | 3.4 | $6.5702 \times 10^3$ | 3.5 | -0.1 | -1.05 |
| $S_{>.4}$ | $1.7877 \times 10^3$ | 1.8 | $1.7945 \times 10^3$ | 1.8 | -0.4 | -2.67 |

**Table 18:** *s*_1_ = MS, *s*_2_ = COSI2; model: bottleneck

| feature | $s_1$ mean | $s_1$ s.e. | $s_2$ mean | $s_2$ s.e. | $\Delta\%$ | z-score |
| --- | --- | --- | --- | --- | --- | --- |
| $r_{50}$ | $1.4335 \times 10^{-1}$ | $1.0 \times 10^{-4}$ | $1.4333 \times 10^{-1}$ | $1.0 \times 10^{-4}$ | 0.0 | 0.14 |
| $r_{100}$ | $9.5984 \times 10^{-2}$ | $7.7 \times 10^{-5}$ | $9.5974 \times 10^{-2}$ | $8.0 \times 10^{-5}$ | 0.0 | 0.09 |
| $r_{150}$ | $8.5805 \times 10^{-2}$ | $7.1 \times 10^{-5}$ | $8.5847 \times 10^{-2}$ | $7.4 \times 10^{-5}$ | -0.0 | -0.41 |
| $r_{50}^2$ | $4.3746 \times 10^{-2}$ | $9.9 \times 10^{-5}$ | $4.3569 \times 10^{-2}$ | $9.8 \times 10^{-5}$ | 0.4 | 1.27 |
| $r_{100}^2$ | $1.1788 \times 10^{-2}$ | $6.7 \times 10^{-5}$ | $1.1653 \times 10^{-2}$ | $6.5 \times 10^{-5}$ | 1.2 | 1.45 |
| $r_{150}^2$ | $8.3878 \times 10^{-3}$ | $5.7 \times 10^{-5}$ | $8.2813 \times 10^{-3}$ | $5.6 \times 10^{-5}$ | 1.3 | 1.33 |
| $\theta$ | $7.4371 \times 10^{-4}$ | $3.6 \times 10^{-7}$ | $7.4432 \times 10^{-4}$ | $3.6 \times 10^{-7}$ | -0.1 | -1.20 |
| S | $1.3956 \times 10^4$ | 6.9 | $1.3962 \times 10^4$ | 6.8 | -0.0 | -0.62 |
| $S_{<.1}$ | $5.7377 \times 10^3$ | 5.2 | $5.7341 \times 10^3$ | 5.2 | 0.1 | 0.49 |
| $S_{\in[.1,.4]}$ | $6.4319 \times 10^3$ | 3.9 | $6.4396 \times 10^3$ | 4.0 | -0.1 | -1.38 |
| $S_{>.4}$ | $1.7860 \times 10^3$ | 1.8 | $1.7883 \times 10^3$ | 1.9 | -0.1 | -0.88 |

**Table 19:** *s*_1_ = MS, *s*_2_ = COSI2; model: island model

| feature | $s_1$ mean | $s_1$ s.e. | $s_2$ mean | $s_2$ s.e. | $\Delta\%$ | z-score |
| --- | --- | --- | --- | --- | --- | --- |
| $r_{50}$ | $7.1124 \times 10^{-2}$ | $2.8 \times 10^{-5}$ | $7.1052 \times 10^{-2}$ | $2.8 \times 10^{-5}$ | 0.1 | 1.82 |
| $r_{100}$ | $4.1675 \times 10^{-2}$ | $1.6 \times 10^{-5}$ | $4.1665 \times 10^{-2}$ | $1.5 \times 10^{-5}$ | 0.0 | 0.46 |
| $r_{150}$ | $3.6240 \times 10^{-2}$ | $1.3 \times 10^{-5}$ | $3.6246 \times 10^{-2}$ | $1.3 \times 10^{-5}$ | -0.0 | -0.33 |
| $r_{50}^2$ | $1.8496 \times 10^{-2}$ | $2.9 \times 10^{-5}$ | $1.8388 \times 10^{-2}$ | $2.9 \times 10^{-5}$ | 0.6 | 2.63 |
| $r_{100}^2$ | $2.7469 \times 10^{-3}$ | $1.3 \times 10^{-5}$ | $2.7295 \times 10^{-3}$ | $1.4 \times 10^{-5}$ | 0.6 | 0.91 |
| $r_{150}^2$ | $1.4771 \times 10^{-3}$ | $1.1 \times 10^{-5}$ | $1.4673 \times 10^{-3}$ | $1.1 \times 10^{-5}$ | 0.7 | 0.63 |
| $\theta$ | $8.1444 \times 10^{-4}$ | $3.4 \times 10^{-7}$ | $8.1551 \times 10^{-4}$ | $3.4 \times 10^{-7}$ | -0.1 | -2.23 |
| S | $2.0163 \times 10^4$ | 6.4 | $2.0183 \times 10^4$ | 6.3 | -0.1 | -2.23 |
| $S_{<.1}$ | $1.1493 \times 10^4$ | 4.7 | $1.1500 \times 10^4$ | 4.6 | -0.1 | -1.06 |
| $S_{\in[.1,.4]}$ | $6.9126 \times 10^3$ | 3.6 | $6.9251 \times 10^3$ | 3.6 | -0.2 | -2.46 |
| $S_{>.4}$ | $1.7576 \times 10^3$ | 1.8 | $1.7579 \times 10^3$ | 1.8 | -0.0 | -0.12 |

**Table 20:** *s*_1_ = MS, *s*_2_ = COSI2; model: island model, high migration

| feature | $s_1$ mean | $s_1$ s.e. | $s_2$ mean | $s_2$ s.e. | $\Delta\%$ | z-score |
| --- | --- | --- | --- | --- | --- | --- |
| $r_{50}$ | $7.1115 \times 10^{-2}$ | $2.8 \times 10^{-5}$ | $7.1094 \times 10^{-2}$ | $2.8 \times 10^{-5}$ | 0.0 | 0.53 |
| $r_{100}$ | $4.1623 \times 10^{-2}$ | $1.5 \times 10^{-5}$ | $4.1603 \times 10^{-2}$ | $1.5 \times 10^{-5}$ | 0.0 | 0.94 |
| $r_{150}$ | $3.6161 \times 10^{-2}$ | $1.3 \times 10^{-5}$ | $3.6156 \times 10^{-2}$ | $1.3 \times 10^{-5}$ | 0.0 | 0.27 |
| $r_{50}^2$ | $1.8346 \times 10^{-2}$ | $2.8 \times 10^{-5}$ | $1.8367 \times 10^{-2}$ | $2.8 \times 10^{-5}$ | -0.1 | -0.53 |
| $r_{100}^2$ | $2.6739 \times 10^{-3}$ | $1.3 \times 10^{-5}$ | $2.6820 \times 10^{-3}$ | $1.3 \times 10^{-5}$ | -0.3 | -0.44 |
| $r_{150}^2$ | $1.4173 \times 10^{-3}$ | $1.1 \times 10^{-5}$ | $1.4287 \times 10^{-3}$ | $1.1 \times 10^{-5}$ | -0.8 | -0.73 |
| $\theta$ | $8.0187 \times 10^{-4}$ | $3.4 \times 10^{-7}$ | $8.0114 \times 10^{-4}$ | $3.4 \times 10^{-7}$ | 0.1 | 1.52 |
| S | $1.9848 \times 10^4$ | 6.1 | $1.9842 \times 10^4$ | 6.3 | 0.0 | 0.68 |
| $S_{<.1}$ | $1.1310 \times 10^4$ | 4.6 | $1.1316 \times 10^4$ | 4.6 | -0.1 | -0.92 |
| $S_{\in[.1,.4]}$ | $6.8066 \times 10^3$ | 3.5 | $6.7946 \times 10^3$ | 3.6 | 0.2 | 2.39 |
| $S_{>.4}$ | $1.7310 \times 10^3$ | 1.7 | $1.7314 \times 10^3$ | 1.8 | -0.0 | -0.16 |

**Table 21:**
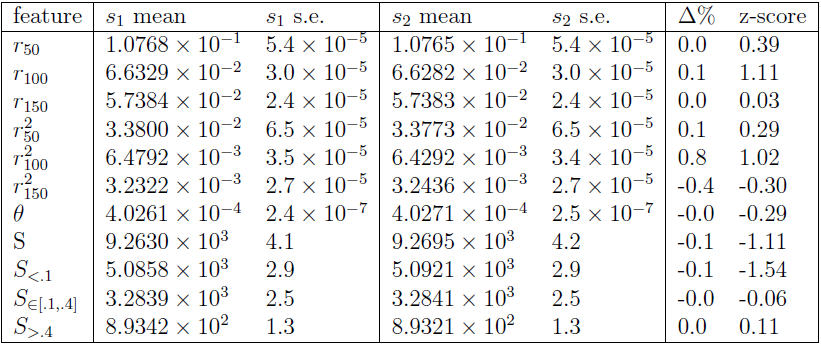
*s*_1_ = MS, *s*_2_ = COSI2; model: piecewise model

| feature | $s_1$ mean | $s_1$ s.e. | $s_2$ mean | $s_2$ s.e. | $\Delta\%$ | z-score |
| --- | --- | --- | --- | --- | --- | --- |
| $r_{50}$ | $1.0768 \times 10^{-1}$ | $5.4 \times 10^{-5}$ | $1.0765 \times 10^{-1}$ | $5.4 \times 10^{-5}$ | 0.0 | 0.39 |
| $r_{100}$ | $6.6329 \times 10^{-2}$ | $3.0 \times 10^{-5}$ | $6.6282 \times 10^{-2}$ | $3.0 \times 10^{-5}$ | 0.1 | 1.11 |
| $r_{150}$ | $5.7384 \times 10^{-2}$ | $2.4 \times 10^{-5}$ | $5.7383 \times 10^{-2}$ | $2.4 \times 10^{-5}$ | 0.0 | 0.03 |
| $r_{50}^2$ | $3.3800 \times 10^{-2}$ | $6.5 \times 10^{-5}$ | $3.3773 \times 10^{-2}$ | $6.5 \times 10^{-5}$ | 0.1 | 0.29 |
| $r_{100}^2$ | $6.4792 \times 10^{-3}$ | $3.5 \times 10^{-5}$ | $6.4292 \times 10^{-3}$ | $3.4 \times 10^{-5}$ | 0.8 | 1.02 |
| $r_{150}^2$ | $3.2322 \times 10^{-3}$ | $2.7 \times 10^{-5}$ | $3.2436 \times 10^{-3}$ | $2.7 \times 10^{-5}$ | -0.4 | -0.30 |
| $\theta$ | $4.0261 \times 10^{-4}$ | $2.4 \times 10^{-7}$ | $4.0271 \times 10^{-4}$ | $2.5 \times 10^{-7}$ | -0.0 | -0.29 |
| S | $9.2630 \times 10^3$ | 4.1 | $9.2695 \times 10^3$ | 4.2 | -0.1 | -1.11 |
| $S_{<.1}$ | $5.0858 \times 10^3$ | 2.9 | $5.0921 \times 10^3$ | 2.9 | -0.1 | -1.54 |
| $S_{\in[.1,.4]}$ | $3.2839 \times 10^3$ | 2.5 | $3.2841 \times 10^3$ | 2.5 | -0.0 | -0.06 |
| $S_{>.4}$ | $8.9342 \times 10^2$ | 1.3 | $8.9321 \times 10^2$ | 1.3 | 0.0 | 0.11 |

#### 3.4 MSprime and COSI2

**Table 22:** *s*_1_ = MSprime, *s*_2_ = COSI2; model: constant size

| feature | $s_1$ mean | $s_1$ s.e. | $s_2$ mean | $s_2$ s.e. | $\Delta\%$ | z-score |
| --- | --- | --- | --- | --- | --- | --- |
| $r_{50}$ | $9.2136 \times 10^{-2}$ | $3.5 \times 10^{-5}$ | $9.2133 \times 10^{-2}$ | $3.5 \times 10^{-5}$ | 0.0 | 0.06 |
| $r_{100}$ | $5.8547 \times 10^{-2}$ | $2.1 \times 10^{-5}$ | $5.8514 \times 10^{-2}$ | $2.1 \times 10^{-5}$ | 0.1 | 1.11 |
| $r_{150}$ | $5.2586 \times 10^{-2}$ | $1.8 \times 10^{-5}$ | $5.2567 \times 10^{-2}$ | $1.9 \times 10^{-5}$ | 0.0 | 0.73 |
| $r_{50}^2$ | $2.2021 \times 10^{-2}$ | $3.6 \times 10^{-5}$ | $2.2038 \times 10^{-2}$ | $3.6 \times 10^{-5}$ | -0.1 | -0.33 |
| $r_{100}^2$ | $3.1562 \times 10^{-3}$ | $1.8 \times 10^{-5}$ | $3.1749 \times 10^{-3}$ | $1.8 \times 10^{-5}$ | -0.6 | -0.73 |
| $r_{150}^2$ | $1.6598 \times 10^{-3}$ | $1.6 \times 10^{-5}$ | $1.6671 \times 10^{-3}$ | $1.5 \times 10^{-5}$ | -0.4 | -0.33 |
| $\theta$ | $7.9935 \times 10^{-4}$ | $3.5 \times 10^{-7}$ | $7.9991 \times 10^{-4}$ | $3.4 \times 10^{-7}$ | -0.1 | -1.15 |
| S | $1.7909 \times 10^4$ | 6.3 | $1.7922 \times 10^4$ | 6.0 | -0.1 | -1.49 |
| $S_{<.1}$ | $9.5580 \times 10^3$ | 4.5 | $9.5658 \times 10^3$ | 4.3 | -0.1 | -1.25 |
| $S_{\in[.1,.4]}$ | $6.5665 \times 10^3$ | 3.5 | $6.5698 \times 10^3$ | 3.5 | -0.1 | -0.67 |
| $S_{>.4}$ | $1.7847 \times 10^3$ | 1.8 | $1.7868 \times 10^3$ | 1.8 | -0.1 | -0.82 |

**Table 23:** *s*_1_ = MSprime, *s*_2_ = COSI2; model: exponential expansion

| feature | $s_1$ mean | $s_1$ s.e. | $s_2$ mean | $s_2$ s.e. | $\Delta\%$ | z-score |
| --- | --- | --- | --- | --- | --- | --- |
| $r_{50}$ | $8.1223 \times 10^{-2}$ | $2.4 \times 10^{-5}$ | $8.1219 \times 10^{-2}$ | $2.4 \times 10^{-5}$ | 0.0 | 0.12 |
| $r_{100}$ | $5.1483 \times 10^{-2}$ | $1.2 \times 10^{-5}$ | $5.1497 \times 10^{-2}$ | $1.2 \times 10^{-5}$ | -0.0 | -0.82 |
| $r_{150}$ | $4.6682 \times 10^{-2}$ | $9.6 \times 10^{-6}$ | $4.6671 \times 10^{-2}$ | $9.6 \times 10^{-6}$ | 0.0 | 0.81 |
| $r_{50}^2$ | $1.8091 \times 10^{-2}$ | $2.8 \times 10^{-5}$ | $1.8078 \times 10^{-2}$ | $2.8 \times 10^{-5}$ | 0.1 | 0.33 |
| $r_{100}^2$ | $2.0507 \times 10^{-3}$ | $1.3 \times 10^{-5}$ | $2.0556 \times 10^{-3}$ | $1.3 \times 10^{-5}$ | -0.2 | -0.27 |
| $r_{150}^2$ | $8.8355 \times 10^{-4}$ | $1.1 \times 10^{-5}$ | $8.6612 \times 10^{-4}$ | $1.2 \times 10^{-5}$ | 2.0 | 1.07 |
| $\theta$ | $8.1226 \times 10^{-4}$ | $3.4 \times 10^{-7}$ | $8.1274 \times 10^{-4}$ | $3.4 \times 10^{-7}$ | -0.1 | -1.00 |
| S | $1.9368 \times 10^4$ | 5.9 | $1.9373 \times 10^4$ | 5.8 | -0.0 | -0.60 |
| $S_{<.1}$ | $1.1009 \times 10^4$ | 4.1 | $1.1008 \times 10^4$ | 3.9 | 0.0 | 0.18 |
| $S_{\in[.1,.4]}$ | $6.5694 \times 10^3$ | 3.4 | $6.5702 \times 10^3$ | 3.5 | -0.0 | -0.16 |
| $S_{>.4}$ | $1.7889 \times 10^3$ | 1.8 | $1.7945 \times 10^3$ | 1.8 | -0.3 | -2.20 |

**Table 24:**
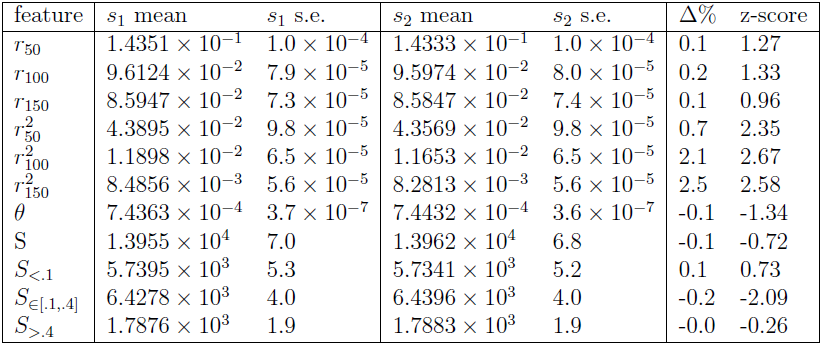
*s*_1_ = MSprime, *s*_2_ = COSI2; model: bottleneck

| feature | $s_1$ mean | $s_1$ s.e. | $s_2$ mean | $s_2$ s.e. | $\Delta\%$ | z-score |
| --- | --- | --- | --- | --- | --- | --- |
| $r_{50}$ | $1.4351 \times 10^{-1}$ | $1.0 \times 10^{-4}$ | $1.4333 \times 10^{-1}$ | $1.0 \times 10^{-4}$ | 0.1 | 1.27 |
| $r_{100}$ | $9.6124 \times 10^{-2}$ | $7.9 \times 10^{-5}$ | $9.5974 \times 10^{-2}$ | $8.0 \times 10^{-5}$ | 0.2 | 1.33 |
| $r_{150}$ | $8.5947 \times 10^{-2}$ | $7.3 \times 10^{-5}$ | $8.5847 \times 10^{-2}$ | $7.4 \times 10^{-5}$ | 0.1 | 0.96 |
| $r_{50}^2$ | $4.3895 \times 10^{-2}$ | $9.8 \times 10^{-5}$ | $4.3569 \times 10^{-2}$ | $9.8 \times 10^{-5}$ | 0.7 | 2.35 |
| $r_{100}^2$ | $1.1898 \times 10^{-2}$ | $6.5 \times 10^{-5}$ | $1.1653 \times 10^{-2}$ | $6.5 \times 10^{-5}$ | 2.1 | 2.67 |
| $r_{150}^2$ | $8.4856 \times 10^{-3}$ | $5.6 \times 10^{-5}$ | $8.2813 \times 10^{-3}$ | $5.6 \times 10^{-5}$ | 2.5 | 2.58 |
| $\theta$ | $7.4363 \times 10^{-4}$ | $3.7 \times 10^{-7}$ | $7.4432 \times 10^{-4}$ | $3.6 \times 10^{-7}$ | -0.1 | -1.34 |
| S | $1.3955 \times 10^4$ | 7.0 | $1.3962 \times 10^4$ | 6.8 | -0.1 | -0.72 |
| $S_{<.1}$ | $5.7395 \times 10^3$ | 5.3 | $5.7341 \times 10^3$ | 5.2 | 0.1 | 0.73 |
| $S_{\in[.1,.4]}$ | $6.4278 \times 10^3$ | 4.0 | $6.4396 \times 10^3$ | 4.0 | -0.2 | -2.09 |
| $S_{>.4}$ | $1.7876 \times 10^3$ | 1.9 | $1.7883 \times 10^3$ | 1.9 | -0.0 | -0.26 |

**Table 25:** *s*_1_ = MSprime, *s*_2_ = COSI2; model: piecewise model

| feature | $s_1$ mean | $s_1$ s.e. | $s_2$ mean | $s_2$ s.e. | $\Delta\%$ | z-score |
| --- | --- | --- | --- | --- | --- | --- |
| $r_{50}$ | $1.0780 \times 10^{-1}$ | $5.5 \times 10^{-5}$ | $1.0765 \times 10^{-1}$ | $5.4 \times 10^{-5}$ | 0.1 | 1.95 |
| $r_{100}$ | $6.6363 \times 10^{-2}$ | $3.0 \times 10^{-5}$ | $6.6282 \times 10^{-2}$ | $3.0 \times 10^{-5}$ | 0.1 | 1.91 |
| $r_{150}$ | $5.7390 \times 10^{-2}$ | $2.4 \times 10^{-5}$ | $5.7383 \times 10^{-2}$ | $2.4 \times 10^{-5}$ | 0.0 | 0.21 |
| $r_{50}^2$ | $3.3855 \times 10^{-2}$ | $6.5 \times 10^{-5}$ | $3.3773 \times 10^{-2}$ | $6.5 \times 10^{-5}$ | 0.2 | 0.89 |
| $r_{100}^2$ | $6.4790 \times 10^{-3}$ | $3.4 \times 10^{-5}$ | $6.4292 \times 10^{-3}$ | $3.4 \times 10^{-5}$ | 0.8 | 1.04 |
| $r_{150}^2$ | $3.2605 \times 10^{-3}$ | $2.7 \times 10^{-5}$ | $3.2436 \times 10^{-3}$ | $2.7 \times 10^{-5}$ | 0.5 | 0.44 |
| $\theta$ | $4.0280 \times 10^{-4}$ | $2.4 \times 10^{-7}$ | $4.0271 \times 10^{-4}$ | $2.5 \times 10^{-7}$ | 0.0 | 0.26 |
| S | $9.2621 \times 10^3$ | 4.2 | $9.2695 \times 10^3$ | 4.2 | -0.1 | -1.25 |
| $S_{<.1}$ | $5.0827 \times 10^3$ | 2.9 | $5.0921 \times 10^3$ | 2.9 | -0.2 | -2.29 |
| $S_{\in[.1,.4]}$ | $3.2823 \times 10^3$ | 2.5 | $3.2841 \times 10^3$ | 2.5 | -0.1 | -0.51 |
| $S_{>.4}$ | $8.9714 \times 10^2$ | 1.3 | $8.9321 \times 10^2$ | 1.3 | 0.4 | 2.14 |

#### 3.5 MSprime and ARGON

**Table 26:** *s*_1_ = MSprime, *s*_2_ = ARGON; model: constant size

| feature | $s_1$ mean | $s_1$ s.e. | $s_2$ mean | $s_2$ s.e. | $\Delta\%$ | z-score |
| --- | --- | --- | --- | --- | --- | --- |
| $r_{50}$ | $9.2136 \times 10^{-2}$ | $3.5 \times 10^{-5}$ | $9.2195 \times 10^{-2}$ | $3.5 \times 10^{-5}$ | -0.1 | -1.19 |
| $r_{100}$ | $5.8547 \times 10^{-2}$ | $2.1 \times 10^{-5}$ | $5.8528 \times 10^{-2}$ | $2.1 \times 10^{-5}$ | 0.0 | 0.64 |
| $r_{150}$ | $5.2586 \times 10^{-2}$ | $1.8 \times 10^{-5}$ | $5.2589 \times 10^{-2}$ | $1.9 \times 10^{-5}$ | -0.0 | -0.11 |
| $r_{50}^2$ | $2.2021 \times 10^{-2}$ | $3.6 \times 10^{-5}$ | $2.2082 \times 10^{-2}$ | $3.6 \times 10^{-5}$ | -0.3 | -1.20 |
| $r_{100}^2$ | $3.1562 \times 10^{-3}$ | $1.8 \times 10^{-5}$ | $3.1710 \times 10^{-3}$ | $1.8 \times 10^{-5}$ | -0.5 | -0.58 |
| $r_{150}^2$ | $1.6598 \times 10^{-3}$ | $1.6 \times 10^{-5}$ | $1.6683 \times 10^{-3}$ | $1.5 \times 10^{-5}$ | -0.5 | -0.39 |
| $\theta$ | $7.9935 \times 10^{-4}$ | $3.5 \times 10^{-7}$ | $7.9992 \times 10^{-4}$ | $3.4 \times 10^{-7}$ | -0.1 | -1.17 |
| S | $1.7909 \times 10^4$ | 6.3 | $1.7911 \times 10^4$ | 6.1 | -0.0 | -0.23 |
| $S_{<.1}$ | $9.5580 \times 10^3$ | 4.5 | $9.5553 \times 10^3$ | 4.4 | 0.0 | 0.43 |
| $S_{\in[.1,.4]}$ | $6.5665 \times 10^3$ | 3.5 | $6.5648 \times 10^3$ | 3.5 | 0.0 | 0.34 |
| $S_{>.4}$ | $1.7847 \times 10^3$ | 1.8 | $1.7905 \times 10^3$ | 1.8 | -0.3 | -2.28 |

**Table 27:** *s*_1_ = MSprime, *s*_2_ = ARGON; model: exponential expansion

| feature | $s_1$ mean | $s_1$ s.e. | $s_2$ mean | $s_2$ s.e. | $\Delta\%$ | z-score |
| --- | --- | --- | --- | --- | --- | --- |
| $r_{50}$ | $8.1223 \times 10^{-2}$ | $2.4 \times 10^{-5}$ | $8.1185 \times 10^{-2}$ | $2.4 \times 10^{-5}$ | 0.0 | 1.12 |
| $r_{100}$ | $5.1483 \times 10^{-2}$ | $1.2 \times 10^{-5}$ | $5.1489 \times 10^{-2}$ | $1.2 \times 10^{-5}$ | -0.0 | -0.35 |
| $r_{150}$ | $4.6682 \times 10^{-2}$ | $9.6 \times 10^{-6}$ | $4.6677 \times 10^{-2}$ | $9.5 \times 10^{-6}$ | 0.0 | 0.37 |
| $r_{50}^2$ | $1.8091 \times 10^{-2}$ | $2.8 \times 10^{-5}$ | $1.8069 \times 10^{-2}$ | $2.8 \times 10^{-5}$ | 0.1 | 0.56 |
| $r_{100}^2$ | $2.0507 \times 10^{-3}$ | $1.3 \times 10^{-5}$ | $2.0460 \times 10^{-3}$ | $1.3 \times 10^{-5}$ | 0.2 | 0.26 |
| $r_{150}^2$ | $8.8355 \times 10^{-4}$ | $1.1 \times 10^{-5}$ | $8.6980 \times 10^{-4}$ | $1.1 \times 10^{-5}$ | 1.6 | 0.88 |
| $\theta$ | $8.1226 \times 10^{-4}$ | $3.4 \times 10^{-7}$ | $8.1197 \times 10^{-4}$ | $3.4 \times 10^{-7}$ | 0.0 | 0.60 |
| S | $1.9368 \times 10^4$ | 5.9 | $1.9363 \times 10^4$ | 5.8 | 0.0 | 0.60 |
| $S_{<.1}$ | $1.1009 \times 10^4$ | 4.1 | $1.1007 \times 10^4$ | 4.0 | 0.0 | 0.35 |
| $S_{\in[.1,.4]}$ | $6.5694 \times 10^3$ | 3.4 | $6.5682 \times 10^3$ | 3.4 | 0.0 | 0.25 |
| $S_{>.4}$ | $1.7889 \times 10^3$ | 1.8 | $1.7880 \times 10^3$ | 1.8 | 0.1 | 0.35 |

**Table 28:** *s*_1_ = MSprime, *s*_2_ = ARGON; model: bottleneck

| feature | $s_1$ mean | $s_1$ s.e. | $s_2$ mean | $s_2$ s.e. | $\Delta\%$ | z-score |
| --- | --- | --- | --- | --- | --- | --- |
| $r_{50}$ | $1.4351 \times 10^{-1}$ | $1.0 \times 10^{-4}$ | $1.4323 \times 10^{-1}$ | $1.0 \times 10^{-4}$ | 0.2 | 1.98 |
| $r_{100}$ | $9.6124 \times 10^{-2}$ | $7.9 \times 10^{-5}$ | $9.5918 \times 10^{-2}$ | $7.7 \times 10^{-5}$ | 0.2 | 1.87 |
| $r_{150}$ | $8.5947 \times 10^{-2}$ | $7.3 \times 10^{-5}$ | $8.5765 \times 10^{-2}$ | $7.2 \times 10^{-5}$ | 0.2 | 1.78 |
| $r_{50}^2$ | $4.3895 \times 10^{-2}$ | $9.8 \times 10^{-5}$ | $4.3686 \times 10^{-2}$ | $1.0 \times 10^{-4}$ | 0.5 | 1.49 |
| $r_{100}^2$ | $1.1898 \times 10^{-2}$ | $6.5 \times 10^{-5}$ | $1.1734 \times 10^{-2}$ | $6.4 \times 10^{-5}$ | 1.4 | 1.80 |
| $r_{150}^2$ | $8.4856 \times 10^{-3}$ | $5.6 \times 10^{-5}$ | $8.3447 \times 10^{-3}$ | $5.6 \times 10^{-5}$ | 1.7 | 1.78 |
| $\theta$ | $7.4363 \times 10^{-4}$ | $3.7 \times 10^{-7}$ | $7.4407 \times 10^{-4}$ | $3.7 \times 10^{-7}$ | -0.1 | -0.84 |
| S | $1.3955 \times 10^4$ | 7.0 | $1.3966 \times 10^4$ | 6.8 | -0.1 | -1.13 |
| $S_{<.1}$ | $5.7395 \times 10^3$ | 5.3 | $5.7458 \times 10^3$ | 5.1 | -0.1 | -0.86 |
| $S_{\in[.1,.4]}$ | $6.4278 \times 10^3$ | 4.0 | $6.4321 \times 10^3$ | 4.0 | -0.1 | -0.76 |
| $S_{>.4}$ | $1.7876 \times 10^3$ | 1.9 | $1.7883 \times 10^3$ | 1.9 | -0.0 | -0.26 |

**Table 29:**
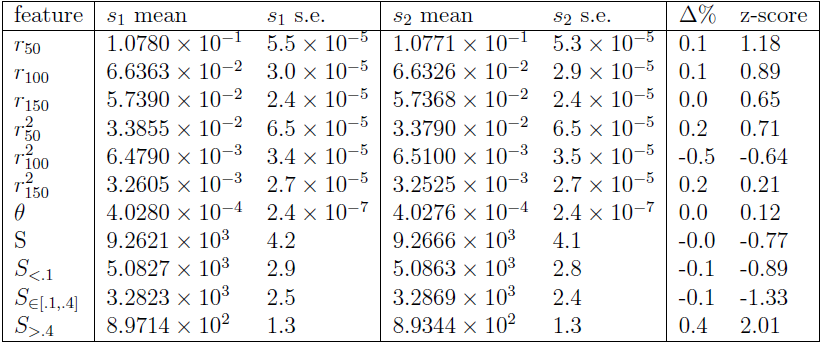
*s*_1_ = MSprime, *s*_2_ = ARGON; model: piecewise model

#### 3.6 MSprime and MS

**Table 30:** *s*_1_ = MSprime, *s*_2_ = MS; model: constant size

| feature | $s_1$ mean | $s_1$ s.e. | $s_2$ mean | $s_2$ s.e. | $\Delta\%$ | z-score |
| --- | --- | --- | --- | --- | --- | --- |
| $r_{50}$ | $9.2136 \times 10^{-2}$ | $3.5 \times 10^{-5}$ | $9.2157 \times 10^{-2}$ | $3.5 \times 10^{-5}$ | -0.0 | -0.42 |
| $r_{100}$ | $5.8547 \times 10^{-2}$ | $2.1 \times 10^{-5}$ | $5.8524 \times 10^{-2}$ | $2.1 \times 10^{-5}$ | 0.0 | 0.77 |
| $r_{150}$ | $5.2586 \times 10^{-2}$ | $1.8 \times 10^{-5}$ | $5.2587 \times 10^{-2}$ | $1.8 \times 10^{-5}$ | -0.0 | -0.04 |
| $r_{50}^2$ | $2.2021 \times 10^{-2}$ | $3.6 \times 10^{-5}$ | $2.2059 \times 10^{-2}$ | $3.6 \times 10^{-5}$ | -0.2 | -0.75 |
| $r_{100}^2$ | $3.1562 \times 10^{-3}$ | $1.8 \times 10^{-5}$ | $3.1556 \times 10^{-3}$ | $1.8 \times 10^{-5}$ | 0.0 | 0.02 |
| $r_{150}^2$ | $1.6598 \times 10^{-3}$ | $1.6 \times 10^{-5}$ | $1.6424 \times 10^{-3}$ | $1.6 \times 10^{-5}$ | 1.1 | 0.77 |
| $\theta$ | $7.9935 \times 10^{-4}$ | $3.5 \times 10^{-7}$ | $7.9966 \times 10^{-4}$ | $3.4 \times 10^{-7}$ | -0.0 | -0.64 |
| S | $1.7909 \times 10^4$ | 6.3 | $1.7916 \times 10^4$ | 6.1 | -0.0 | -0.80 |
| $S_{<.1}$ | $9.5580 \times 10^3$ | 4.5 | $9.5639 \times 10^3$ | 4.4 | -0.1 | -0.94 |
| $S_{\in[.1,.4]}$ | $6.5665 \times 10^3$ | 3.5 | $6.5642 \times 10^3$ | 3.4 | 0.0 | 0.47 |
| $S_{>.4}$ | $1.7847 \times 10^3$ | 1.8 | $1.7879 \times 10^3$ | 1.8 | -0.2 | -1.26 |

**Table 31:** *s*_1_ = MSprime, *s*_2_ = MS; model: exponential expansion

| feature | $s_1$ mean | $s_1$ s.e. | $s_2$ mean | $s_2$ s.e. | $\Delta\%$ | z-score |
| --- | --- | --- | --- | --- | --- | --- |
| $r_{50}$ | $8.1223 \times 10^{-2}$ | $2.4 \times 10^{-5}$ | $8.1150 \times 10^{-2}$ | $2.4 \times 10^{-5}$ | 0.1 | 2.15 |
| $r_{100}$ | $5.1483 \times 10^{-2}$ | $1.2 \times 10^{-5}$ | $5.1469 \times 10^{-2}$ | $1.2 \times 10^{-5}$ | 0.0 | 0.82 |
| $r_{150}$ | $4.6682 \times 10^{-2}$ | $9.6 \times 10^{-6}$ | $4.6659 \times 10^{-2}$ | $9.5 \times 10^{-6}$ | 0.0 | 1.70 |
| $r_{50}^2$ | $1.8091 \times 10^{-2}$ | $2.8 \times 10^{-5}$ | $1.8049 \times 10^{-2}$ | $2.9 \times 10^{-5}$ | 0.2 | 1.04 |
| $r_{100}^2$ | $2.0507 \times 10^{-3}$ | $1.3 \times 10^{-5}$ | $2.0228 \times 10^{-3}$ | $1.3 \times 10^{-5}$ | 1.4 | 1.52 |
| $r_{150}^2$ | $8.8355 \times 10^{-4}$ | $1.1 \times 10^{-5}$ | $8.5858 \times 10^{-4}$ | $1.1 \times 10^{-5}$ | 2.9 | 1.61 |
| $\theta$ | $8.1226 \times 10^{-4}$ | $3.4 \times 10^{-7}$ | $8.1180 \times 10^{-4}$ | $3.4 \times 10^{-7}$ | 0.1 | 0.96 |
| S | $1.9368 \times 10^4$ | 5.9 | $1.9370 \times 10^4$ | 5.8 | -0.0 | -0.24 |
| $S_{<.1}$ | $1.1009 \times 10^4$ | 4.1 | $1.1017 \times 10^4$ | 4.0 | -0.1 | -1.40 |
| $S_{\in[.1,.4]}$ | $6.5694 \times 10^3$ | 3.4 | $6.5651 \times 10^3$ | 3.4 | 0.1 | 0.89 |
| $S_{>.4}$ | $1.7889 \times 10^3$ | 1.8 | $1.7877 \times 10^3$ | 1.8 | 0.1 | 0.47 |

**Table 32:** *s*_1_ = MSprime, *s*_2_ = MS; model: bottleneck

| feature | $s_1$ mean | $s_1$ s.e. | $s_2$ mean | $s_2$ s.e. | $\Delta\%$ | z-score |
| --- | --- | --- | --- | --- | --- | --- |
| $r_{50}$ | $1.4351 \times 10^{-1}$ | $1.0 \times 10^{-4}$ | $1.4335 \times 10^{-1}$ | $1.0 \times 10^{-4}$ | 0.1 | 1.13 |
| $r_{100}$ | $9.6124 \times 10^{-2}$ | $7.9 \times 10^{-5}$ | $9.5984 \times 10^{-2}$ | $7.7 \times 10^{-5}$ | 0.1 | 1.27 |
| $r_{150}$ | $8.5947 \times 10^{-2}$ | $7.3 \times 10^{-5}$ | $8.5805 \times 10^{-2}$ | $7.1 \times 10^{-5}$ | 0.2 | 1.39 |
| $r_{50}^2$ | $4.3895 \times 10^{-2}$ | $9.8 \times 10^{-5}$ | $4.3746 \times 10^{-2}$ | $9.9 \times 10^{-5}$ | 0.3 | 1.07 |
| $r_{100}^2$ | $1.1898 \times 10^{-2}$ | $6.5 \times 10^{-5}$ | $1.1788 \times 10^{-2}$ | $6.7 \times 10^{-5}$ | 0.9 | 1.18 |
| $r_{150}^2$ | $8.4856 \times 10^{-3}$ | $5.6 \times 10^{-5}$ | $8.3878 \times 10^{-3}$ | $5.7 \times 10^{-5}$ | 1.2 | 1.22 |
| $\theta$ | $7.4363 \times 10^{-4}$ | $3.7 \times 10^{-7}$ | $7.4371 \times 10^{-4}$ | $3.6 \times 10^{-7}$ | -0.0 | -0.15 |
| S | $1.3955 \times 10^4$ | 7.0 | $1.3956 \times 10^4$ | 6.9 | -0.0 | -0.10 |
| $S_{<.1}$ | $5.7395 \times 10^3$ | 5.3 | $5.7377 \times 10^3$ | 5.2 | 0.0 | 0.24 |
| $S_{\in[.1,.4]}$ | $6.4278 \times 10^3$ | 4.0 | $6.4319 \times 10^3$ | 3.9 | -0.1 | -0.73 |
| $S_{>.4}$ | $1.7876 \times 10^3$ | 1.9 | $1.7860 \times 10^3$ | 1.8 | 0.1 | 0.61 |

**Table 33:** *s*_1_ = MSprime, *s*_2_ = MS; model: piecewise model

| feature | $s_1$ mean | $s_1$ s.e. | $s_2$ mean | $s_2$ s.e. | $\Delta\%$ | z-score |
| --- | --- | --- | --- | --- | --- | --- |
| $r_{50}$ | $1.0780 \times 10^{-1}$ | $5.5 \times 10^{-5}$ | $1.0768 \times 10^{-1}$ | $5.4 \times 10^{-5}$ | 0.1 | 1.56 |
| $r_{100}$ | $6.6363 \times 10^{-2}$ | $3.0 \times 10^{-5}$ | $6.6329 \times 10^{-2}$ | $3.0 \times 10^{-5}$ | 0.1 | 0.80 |
| $r_{150}$ | $5.7390 \times 10^{-2}$ | $2.4 \times 10^{-5}$ | $5.7384 \times 10^{-2}$ | $2.4 \times 10^{-5}$ | 0.0 | 0.18 |
| $r_{50}^2$ | $3.3855 \times 10^{-2}$ | $6.5 \times 10^{-5}$ | $3.3800 \times 10^{-2}$ | $6.5 \times 10^{-5}$ | 0.2 | 0.60 |
| $r_{100}^2$ | $6.4790 \times 10^{-3}$ | $3.4 \times 10^{-5}$ | $6.4792 \times 10^{-3}$ | $3.5 \times 10^{-5}$ | -0.0 | -0.00 |
| $r_{150}^2$ | $3.2605 \times 10^{-3}$ | $2.7 \times 10^{-5}$ | $3.2322 \times 10^{-3}$ | $2.7 \times 10^{-5}$ | 0.9 | 0.74 |
| $\theta$ | $4.0280 \times 10^{-4}$ | $2.4 \times 10^{-7}$ | $4.0261 \times 10^{-4}$ | $2.4 \times 10^{-7}$ | 0.0 | 0.56 |
| S | $9.2621 \times 10^3$ | 4.2 | $9.2630 \times 10^3$ | 4.1 | -0.0 | -0.15 |
| $S_{<.1}$ | $5.0827 \times 10^3$ | 2.9 | $5.0858 \times 10^3$ | 2.9 | -0.1 | -0.76 |
| $S_{\in[.1,.4]}$ | $3.2823 \times 10^3$ | 2.5 | $3.2839 \times 10^3$ | 2.5 | -0.0 | -0.45 |
| $S_{>.4}$ | $8.9714 \times 10^2$ | 1.3 | $8.9342 \times 10^2$ | 1.3 | 0.4 | 2.02 |

#### 3.7 SCRM and MS

**Table 34:**
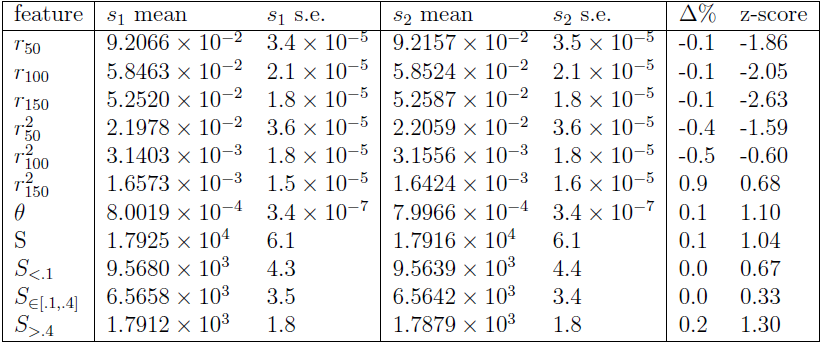
*s*_1_ = SCRM, *s*_2_ = MS; model: constant size

**Table 35:**
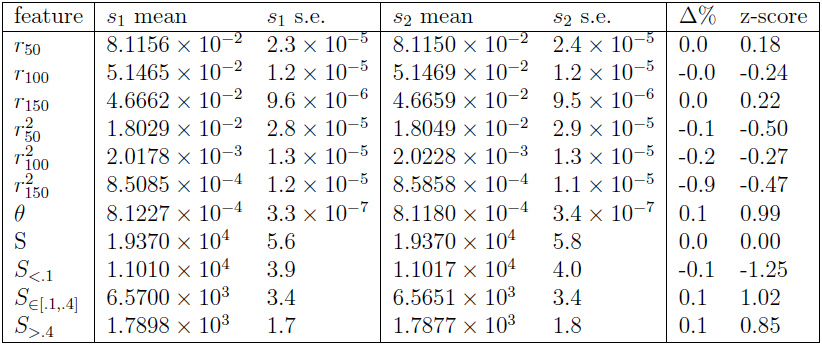
*s*_1_ = SCRM, *s*_2_ = MS; model: exponential expansion

| feature | $s_1$ mean | $s_1$ s.e. | $s_2$ mean | $s_2$ s.e. | $\Delta\%$ | z-score |
| --- | --- | --- | --- | --- | --- | --- |
| $r_{50}$ | $8.1156 \times 10^{-2}$ | $2.3 \times 10^{-5}$ | $8.1150 \times 10^{-2}$ | $2.4 \times 10^{-5}$ | 0.0 | 0.18 |
| $r_{100}$ | $5.1465 \times 10^{-2}$ | $1.2 \times 10^{-5}$ | $5.1469 \times 10^{-2}$ | $1.2 \times 10^{-5}$ | -0.0 | -0.24 |
| $r_{150}$ | $4.6662 \times 10^{-2}$ | $9.6 \times 10^{-6}$ | $4.6659 \times 10^{-2}$ | $9.5 \times 10^{-6}$ | 0.0 | 0.22 |
| $r_{50}^2$ | $1.8029 \times 10^{-2}$ | $2.8 \times 10^{-5}$ | $1.8049 \times 10^{-2}$ | $2.9 \times 10^{-5}$ | -0.1 | -0.50 |
| $r_{100}^2$ | $2.0178 \times 10^{-3}$ | $1.3 \times 10^{-5}$ | $2.0228 \times 10^{-3}$ | $1.3 \times 10^{-5}$ | -0.2 | -0.27 |
| $r_{150}^2$ | $8.5085 \times 10^{-4}$ | $1.2 \times 10^{-5}$ | $8.5858 \times 10^{-4}$ | $1.1 \times 10^{-5}$ | -0.9 | -0.47 |
| $\theta$ | $8.1227 \times 10^{-4}$ | $3.3 \times 10^{-7}$ | $8.1180 \times 10^{-4}$ | $3.4 \times 10^{-7}$ | 0.1 | 0.99 |
| S | $1.9370 \times 10^4$ | 5.6 | $1.9370 \times 10^4$ | 5.8 | 0.0 | 0.00 |
| $S_{<.1}$ | $1.1010 \times 10^4$ | 3.9 | $1.1017 \times 10^4$ | 4.0 | -0.1 | -1.25 |
| $S_{\in[.1,.4]}$ | $6.5700 \times 10^3$ | 3.4 | $6.5651 \times 10^3$ | 3.4 | 0.1 | 1.02 |
| $S_{>.4}$ | $1.7898 \times 10^3$ | 1.7 | $1.7877 \times 10^3$ | 1.8 | 0.1 | 0.85 |

**Table 36:**
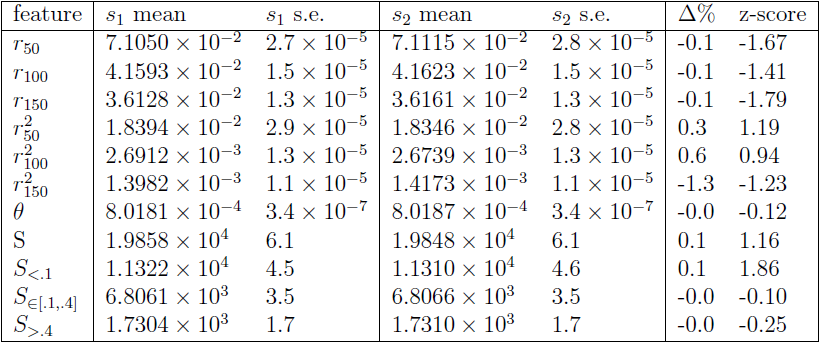
*s*_1_ = SCRM, *s*_2_ = MS; model: island model, high migration

**Table 37:** *s*_1_ = SCRM, *s*_2_ = MS; model: island model

| feature | $s_1$ mean | $s_1$ s.e. | $s_2$ mean | $s_2$ s.e. | $\Delta\%$ | z-score |
| --- | --- | --- | --- | --- | --- | --- |
| $r_{50}$ | $7.1087 \times 10^{-2}$ | $2.7 \times 10^{-5}$ | $7.1124 \times 10^{-2}$ | $2.8 \times 10^{-5}$ | -0.1 | -0.95 |
| $r_{100}$ | $4.1654 \times 10^{-2}$ | $1.5 \times 10^{-5}$ | $4.1675 \times 10^{-2}$ | $1.6 \times 10^{-5}$ | -0.1 | -0.96 |
| $r_{150}$ | $3.6234 \times 10^{-2}$ | $1.3 \times 10^{-5}$ | $3.6240 \times 10^{-2}$ | $1.3 \times 10^{-5}$ | -0.0 | -0.33 |
| $r_{50}^2$ | $1.8399 \times 10^{-2}$ | $2.9 \times 10^{-5}$ | $1.8496 \times 10^{-2}$ | $2.9 \times 10^{-5}$ | -0.5 | -2.37 |
| $r_{100}^2$ | $2.7327 \times 10^{-3}$ | $1.3 \times 10^{-5}$ | $2.7469 \times 10^{-3}$ | $1.3 \times 10^{-5}$ | -0.5 | -0.77 |
| $r_{150}^2$ | $1.4488 \times 10^{-3}$ | $1.1 \times 10^{-5}$ | $1.4771 \times 10^{-3}$ | $1.1 \times 10^{-5}$ | -1.9 | -1.82 |
| $\theta$ | $8.1548 \times 10^{-4}$ | $3.5 \times 10^{-7}$ | $8.1444 \times 10^{-4}$ | $3.4 \times 10^{-7}$ | 0.1 | 2.13 |
| S | $2.0178 \times 10^4$ | 6.2 | $2.0163 \times 10^4$ | 6.4 | 0.1 | 1.68 |
| $S_{<.1}$ | $1.1498 \times 10^4$ | 4.5 | $1.1493 \times 10^4$ | 4.7 | 0.0 | 0.77 |
| $S_{\in[.1,.4]}$ | $6.9192 \times 10^3$ | 3.7 | $6.9126 \times 10^3$ | 3.6 | 0.1 | 1.28 |
| $S_{>.4}$ | $1.7617 \times 10^3$ | 1.8 | $1.7576 \times 10^3$ | 1.8 | 0.2 | 1.61 |

**Table 38:**
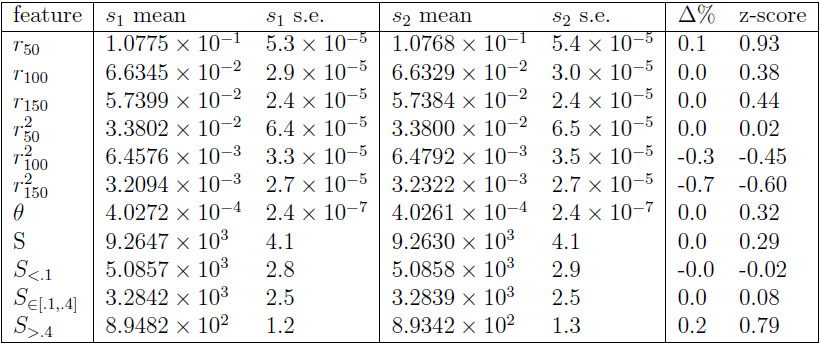
*s*_1_ = SCRM, *s*_2_ = MS; model: piecewise model

| feature | $s_1$ mean | $s_1$ s.e. | $s_2$ mean | $s_2$ s.e. | $\Delta\%$ | z-score |
| --- | --- | --- | --- | --- | --- | --- |
| $r_{50}$ | $1.0775 \times 10^{-1}$ | $5.3 \times 10^{-5}$ | $1.0768 \times 10^{-1}$ | $5.4 \times 10^{-5}$ | 0.1 | 0.93 |
| $r_{100}$ | $6.6345 \times 10^{-2}$ | $2.9 \times 10^{-5}$ | $6.6329 \times 10^{-2}$ | $3.0 \times 10^{-5}$ | 0.0 | 0.38 |
| $r_{150}$ | $5.7399 \times 10^{-2}$ | $2.4 \times 10^{-5}$ | $5.7384 \times 10^{-2}$ | $2.4 \times 10^{-5}$ | 0.0 | 0.44 |
| $r_{50}^2$ | $3.3802 \times 10^{-2}$ | $6.4 \times 10^{-5}$ | $3.3800 \times 10^{-2}$ | $6.5 \times 10^{-5}$ | 0.0 | 0.02 |
| $r_{100}^2$ | $6.4576 \times 10^{-3}$ | $3.3 \times 10^{-5}$ | $6.4792 \times 10^{-3}$ | $3.5 \times 10^{-5}$ | -0.3 | -0.45 |
| $r_{150}^2$ | $3.2094 \times 10^{-3}$ | $2.7 \times 10^{-5}$ | $3.2322 \times 10^{-3}$ | $2.7 \times 10^{-5}$ | -0.7 | -0.60 |
| $\theta$ | $4.0272 \times 10^{-4}$ | $2.4 \times 10^{-7}$ | $4.0261 \times 10^{-4}$ | $2.4 \times 10^{-7}$ | 0.0 | 0.32 |
| S | $9.2647 \times 10^3$ | 4.1 | $9.2630 \times 10^3$ | 4.1 | 0.0 | 0.29 |
| $S_{<.1}$ | $5.0857 \times 10^3$ | 2.8 | $5.0858 \times 10^3$ | 2.9 | -0.0 | -0.02 |
| $S_{\in[.1,.4]}$ | $3.2842 \times 10^3$ | 2.5 | $3.2839 \times 10^3$ | 2.5 | 0.0 | 0.08 |
| $S_{>.4}$ | $8.9482 \times 10^2$ | 1.2 | $8.9342 \times 10^2$ | 1.3 | 0.2 | 0.79 |

#### 3.8 ARGON with approximation and MS

**Table 39:** *s*_1_ = ARGON with 10*μM* approximation, *s*_2_ = MS; model: constant size

| feature | $s_1$ mean | $s_1$ s.e. | $s_2$ mean | $s_2$ s.e. | $\Delta\%$ | z-score |
| --- | --- | --- | --- | --- | --- | --- |
| $r_{50}$ | $9.7980 \times 10^{-2}$ | $4.1 \times 10^{-5}$ | $9.2157 \times 10^{-2}$ | $3.5 \times 10^{-5}$ | 6.3 | 108.02 * |
| $r_{100}$ | $5.8527 \times 10^{-2}$ | $2.2 \times 10^{-5}$ | $5.8524 \times 10^{-2}$ | $2.1 \times 10^{-5}$ | 0.0 | 0.10 |
| $r_{150}$ | $5.2543 \times 10^{-2}$ | $1.9 \times 10^{-5}$ | $5.2587 \times 10^{-2}$ | $1.8 \times 10^{-5}$ | -0.1 | -1.68 |
| $r_{50}^2$ | $2.7954 \times 10^{-2}$ | $4.4 \times 10^{-5}$ | $2.2059 \times 10^{-2}$ | $3.6 \times 10^{-5}$ | 26.7 | 103.69 * |
| $r_{100}^2$ | $3.1551 \times 10^{-3}$ | $2.2 \times 10^{-5}$ | $3.1556 \times 10^{-3}$ | $1.8 \times 10^{-5}$ | -0.0 | -0.02 |
| $r_{150}^2$ | $1.6397 \times 10^{-3}$ | $1.9 \times 10^{-5}$ | $1.6424 \times 10^{-3}$ | $1.6 \times 10^{-5}$ | -0.2 | -0.11 |
| $\theta$ | $8.0000 \times 10^{-4}$ | $3.7 \times 10^{-7}$ | $7.9966 \times 10^{-4}$ | $3.4 \times 10^{-7}$ | 0.0 | 0.68 |
| S | $1.7915 \times 10^4$ | 6.4 | $1.7916 \times 10^4$ | 6.1 | -0.0 | -0.11 |
| $S_{<.1}$ | $9.5557 \times 10^3$ | 4.6 | $9.5639 \times 10^3$ | 4.4 | -0.1 | -1.29 |
| $S_{\in[.1,.4]}$ | $6.5704 \times 10^3$ | 4.0 | $6.5642 \times 10^3$ | 3.4 | 0.1 | 1.18 |
| $S_{>.4}$ | $1.7894 \times 10^3$ | 2.2 | $1.7879 \times 10^3$ | 1.8 | 0.1 | 0.53 |

**Table 40:** *s*_1_ = ARGON with 10*μM* approximation, *s*_2_ = MS; model: exponential expansion

| feature | $s_1$ mean | $s_1$ s.e. | $s_2$ mean | $s_2$ s.e. | $\Delta\%$ | z-score |
| --- | --- | --- | --- | --- | --- | --- |
| $r_{50}$ | $8.6366 \times 10^{-2}$ | $2.9 \times 10^{-5}$ | $8.1150 \times 10^{-2}$ | $2.4 \times 10^{-5}$ | 6.4 | 138.56 * |
| $r_{100}$ | $5.1517 \times 10^{-2}$ | $1.3 \times 10^{-5}$ | $5.1469 \times 10^{-2}$ | $1.2 \times 10^{-5}$ | 0.1 | 2.71 |
| $r_{150}$ | $4.6678 \times 10^{-2}$ | $1.0 \times 10^{-5}$ | $4.6659 \times 10^{-2}$ | $9.5 \times 10^{-6}$ | 0.0 | 1.38 |
| $r_{50}^2$ | $2.3235 \times 10^{-2}$ | $3.5 \times 10^{-5}$ | $1.8049 \times 10^{-2}$ | $2.9 \times 10^{-5}$ | 28.7 | 114.10 * |
| $r_{100}^2$ | $2.0614 \times 10^{-3}$ | $1.6 \times 10^{-5}$ | $2.0228 \times 10^{-3}$ | $1.3 \times 10^{-5}$ | 1.9 | 1.87 |
| $r_{150}^2$ | $8.7392 \times 10^{-4}$ | $1.4 \times 10^{-5}$ | $8.5858 \times 10^{-4}$ | $1.1 \times 10^{-5}$ | 1.8 | 0.86 |
| $\theta$ | $8.1214 \times 10^{-4}$ | $3.7 \times 10^{-7}$ | $8.1180 \times 10^{-4}$ | $3.4 \times 10^{-7}$ | 0.0 | 0.68 |
| S | $1.9366 \times 10^4$ | 6.2 | $1.9370 \times 10^4$ | 5.8 | -0.0 | -0.47 |
| $S_{<.1}$ | $1.1006 \times 10^4$ | 4.2 | $1.1017 \times 10^4$ | 4.0 | -0.1 | -1.90 |
| $S_{\in[.1,.4]}$ | $6.5697 \times 10^3$ | 4.0 | $6.5651 \times 10^3$ | 3.4 | 0.1 | 0.88 |
| $S_{>.4}$ | $1.7897 \times 10^3$ | 2.2 | $1.7877 \times 10^3$ | 1.8 | 0.1 | 0.70 |

**Table 41:** *s*_1_ = ARGON with 50*μM* approximation, *s*_2_ = MS; model: constant size

| feature | $s_1$ mean | $s_1$ s.e. | $s_2$ mean | $s_2$ s.e. | $\Delta\%$ | z-score |
| --- | --- | --- | --- | --- | --- | --- |
| $r_{50}$ | $1.5172 \times 10^{-1}$ | $1.3 \times 10^{-4}$ | $9.2157 \times 10^{-2}$ | $3.5 \times 10^{-5}$ | 64.6 | 442.42 * |
| $r_{100}$ | $5.9439 \times 10^{-2}$ | $3.0 \times 10^{-5}$ | $5.8524 \times 10^{-2}$ | $2.1 \times 10^{-5}$ | 1.6 | 24.99 * |
| $r_{150}$ | $5.2783 \times 10^{-2}$ | $2.5 \times 10^{-5}$ | $5.2587 \times 10^{-2}$ | $1.8 \times 10^{-5}$ | 0.4 | 6.36 * |
| $r_{50}^2$ | $7.8067 \times 10^{-2}$ | $1.5 \times 10^{-4}$ | $2.2059 \times 10^{-2}$ | $3.6 \times 10^{-5}$ | 253.9 | 363.08 * |
| $r_{100}^2$ | $3.5640 \times 10^{-3}$ | $4.6 \times 10^{-5}$ | $3.1556 \times 10^{-3}$ | $1.8 \times 10^{-5}$ | 12.9 | 8.27 * |
| $r_{150}^2$ | $1.7999 \times 10^{-3}$ | $4.2 \times 10^{-5}$ | $1.6424 \times 10^{-3}$ | $1.6 \times 10^{-5}$ | 9.6 | 3.50 * |
| $\theta$ | $8.0020 \times 10^{-4}$ | $5.8 \times 10^{-7}$ | $7.9966 \times 10^{-4}$ | $3.4 \times 10^{-7}$ | 0.1 | 0.80 |
| S | $1.7914 \times 10^4$ | 8.6 | $1.7916 \times 10^4$ | 6.1 | -0.0 | -0.19 |
| $S_{<.1}$ | $9.5523 \times 10^3$ | 5.9 | $9.5639 \times 10^3$ | 4.4 | -0.1 | -1.58 |
| $S_{\in[.1,.4]}$ | $6.5689 \times 10^3$ | 6.6 | $6.5642 \times 10^3$ | 3.4 | 0.1 | 0.63 |
| $S_{>.4}$ | $1.7923 \times 10^3$ | 4.2 | $1.7879 \times 10^3$ | 1.8 | 0.2 | 0.96 |

**Table 42:** *s*_1_ = ARGON with 50*μM* approximation, *s*_2_ = MS; model: exponential expansion

| feature | $s_1$ mean | $s_1$ s.e. | $s_2$ mean | $s_2$ s.e. | $\Delta\%$ | z-score |
| --- | --- | --- | --- | --- | --- | --- |
| $r_{50}$ | $1.3433 \times 10^{-1}$ | $1.2 \times 10^{-4}$ | $8.1150 \times 10^{-2}$ | $2.4 \times 10^{-5}$ | 65.5 | 434.56 * |
| $r_{100}$ | $5.2339 \times 10^{-2}$ | $1.9 \times 10^{-5}$ | $5.1469 \times 10^{-2}$ | $1.2 \times 10^{-5}$ | 1.7 | 38.71 * |
| $r_{150}$ | $4.6874 \times 10^{-2}$ | $1.6 \times 10^{-5}$ | $4.6659 \times 10^{-2}$ | $9.5 \times 10^{-6}$ | 0.5 | 11.55 * |
| $r_{50}^2$ | $6.7000 \times 10^{-2}$ | $1.3 \times 10^{-4}$ | $1.8049 \times 10^{-2}$ | $2.9 \times 10^{-5}$ | 271.2 | 367.51 * |
| $r_{100}^2$ | $2.2971 \times 10^{-3}$ | $3.6 \times 10^{-5}$ | $2.0228 \times 10^{-3}$ | $1.3 \times 10^{-5}$ | 13.6 | 7.17 * |
| $r_{150}^2$ | $9.3514 \times 10^{-4}$ | $3.3 \times 10^{-5}$ | $8.5858 \times 10^{-4}$ | $1.1 \times 10^{-5}$ | 8.9 | 2.20 |
| $\theta$ | $8.1294 \times 10^{-4}$ | $5.9 \times 10^{-7}$ | $8.1180 \times 10^{-4}$ | $3.4 \times 10^{-7}$ | 0.1 | 1.67 |
| S | $1.9377 \times 10^4$ | 8.5 | $1.9370 \times 10^4$ | 5.8 | 0.0 | 0.68 |
| $S_{<.1}$ | $1.1015 \times 10^4$ | 5.6 | $1.1017 \times 10^4$ | 4.0 | -0.0 | -0.29 |
| $S_{\in[.1,.4]}$ | $6.5661 \times 10^3$ | 6.5 | $6.5651 \times 10^3$ | 3.4 | 0.0 | 0.14 |
| $S_{>.4}$ | $1.7962 \times 10^3$ | 4.2 | $1.7877 \times 10^3$ | 1.8 | 0.5 | 1.86 |

#### 3.9 SCRM with approximation and MS

**Table 43:** *s*_1_ = SCRM with “-l” set to 0, *s*_2_ = MS; model: constant size

| feature | $s_1$ mean | $s_1$ s.e. | $s_2$ mean | $s_2$ s.e. | $\Delta\%$ | z-score |
| --- | --- | --- | --- | --- | --- | --- |
| $r_{50}$ | $9.1817 \times 10^{-2}$ | $3.2 \times 10^{-5}$ | $9.2157 \times 10^{-2}$ | $3.5 \times 10^{-5}$ | -0.4 | -7.17 * |
| $r_{100}$ | $5.8418 \times 10^{-2}$ | $2.0 \times 10^{-5}$ | $5.8524 \times 10^{-2}$ | $2.1 \times 10^{-5}$ | -0.2 | -3.66 * |
| $r_{150}$ | $5.2499 \times 10^{-2}$ | $1.8 \times 10^{-5}$ | $5.2587 \times 10^{-2}$ | $1.8 \times 10^{-5}$ | -0.2 | -3.46 * |
| $r_{50}^2$ | $2.1566 \times 10^{-2}$ | $3.3 \times 10^{-5}$ | $2.2059 \times 10^{-2}$ | $3.6 \times 10^{-5}$ | -2.2 | -10.09 * |
| $r_{100}^2$ | $2.9907 \times 10^{-3}$ | $1.8 \times 10^{-5}$ | $3.1556 \times 10^{-3}$ | $1.8 \times 10^{-5}$ | -5.2 | -6.48 * |
| $r_{150}^2$ | $1.5740 \times 10^{-3}$ | $1.5 \times 10^{-5}$ | $1.6424 \times 10^{-3}$ | $1.6 \times 10^{-5}$ | -4.2 | -3.12 |
| $\theta$ | $8.0040 \times 10^{-4}$ | $3.0 \times 10^{-7}$ | $7.9966 \times 10^{-4}$ | $3.4 \times 10^{-7}$ | 0.1 | 1.63 |
| S | $1.7923 \times 10^4$ | 5.1 | $1.7916 \times 10^4$ | 6.1 | 0.0 | 0.88 |
| $S_{<.1}$ | $9.5626 \times 10^3$ | 3.9 | $9.5639 \times 10^3$ | 4.4 | -0.0 | -0.22 |
| $S_{\in[.1,.4]}$ | $6.5705 \times 10^3$ | 3.1 | $6.5642 \times 10^3$ | 3.4 | 0.1 | 1.37 |
| $S_{>.4}$ | $1.7901 \times 10^3$ | 1.7 | $1.7879 \times 10^3$ | 1.8 | 0.1 | 0.89 |

**Table 44:** *s*_1_ = SCRM with “-l” set to 0, *s*_2_ = MS; model: exponential expansion

| feature | $s_1$ mean | $s_1$ s.e. | $s_2$ mean | $s_2$ s.e. | $\Delta\%$ | z-score |
| --- | --- | --- | --- | --- | --- | --- |
| $r_{50}$ | $8.0921 \times 10^{-2}$ | $2.2 \times 10^{-5}$ | $8.1150 \times 10^{-2}$ | $2.4 \times 10^{-5}$ | -0.3 | -7.03 * |
| $r_{100}$ | $5.1395 \times 10^{-2}$ | $1.2 \times 10^{-5}$ | $5.1469 \times 10^{-2}$ | $1.2 \times 10^{-5}$ | -0.1 | -4.36 * |
| $r_{150}$ | $4.6608 \times 10^{-2}$ | $9.6 \times 10^{-6}$ | $4.6659 \times 10^{-2}$ | $9.5 \times 10^{-6}$ | -0.1 | -3.78 * |
| $r_{50}^2$ | $1.7728 \times 10^{-2}$ | $2.6 \times 10^{-5}$ | $1.8049 \times 10^{-2}$ | $2.9 \times 10^{-5}$ | -1.8 | -8.24 * |
| $r_{100}^2$ | $1.9187 \times 10^{-3}$ | $1.3 \times 10^{-5}$ | $2.0228 \times 10^{-3}$ | $1.3 \times 10^{-5}$ | -5.1 | -5.66 * |
| $r_{150}^2$ | $8.0923 \times 10^{-4}$ | $1.1 \times 10^{-5}$ | $8.5858 \times 10^{-4}$ | $1.1 \times 10^{-5}$ | -5.7 | -3.17 |
| $\theta$ | $8.1183 \times 10^{-4}$ | $2.9 \times 10^{-7}$ | $8.1180 \times 10^{-4}$ | $3.4 \times 10^{-7}$ | 0.0 | 0.07 |
| S | $1.9365 \times 10^4$ | 4.7 | $1.9370 \times 10^4$ | 5.8 | -0.0 | -0.67 |
| $S_{<.1}$ | $1.1013 \times 10^4$ | 3.5 | $1.1017 \times 10^4$ | 4.0 | -0.0 | -0.75 |
| $S_{\in[.1,.4]}$ | $6.5611 \times 10^3$ | 3.0 | $6.5651 \times 10^3$ | 3.4 | -0.1 | -0.88 |
| $S_{>.4}$ | $1.7912 \times 10^3$ | 1.7 | $1.7877 \times 10^3$ | 1.8 | 0.2 | 1.41 |

#### 3.10 COSI2 with approximation and MS

**Table 45:** *s*_1_ = COSI2 with “-u” set to 0, *s*_2_ = MS; model: constant size

| feature | $s_1$ mean | $s_1$ s.e. | $s_2$ mean | $s_2$ s.e. | $\Delta\%$ | z-score |
| --- | --- | --- | --- | --- | --- | --- |
| $r_{50}$ | $9.0589 \times 10^{-2}$ | $3.1 \times 10^{-5}$ | $9.2157 \times 10^{-2}$ | $3.5 \times 10^{-5}$ | -1.7 | -33.54 * |
| $r_{100}$ | $5.8253 \times 10^{-2}$ | $2.0 \times 10^{-5}$ | $5.8524 \times 10^{-2}$ | $2.1 \times 10^{-5}$ | -0.5 | -9.34 * |
| $r_{150}$ | $5.2397 \times 10^{-2}$ | $1.8 \times 10^{-5}$ | $5.2587 \times 10^{-2}$ | $1.8 \times 10^{-5}$ | -0.4 | -7.46 * |
| $r_{50}^2$ | $2.0477 \times 10^{-2}$ | $3.2 \times 10^{-5}$ | $2.2059 \times 10^{-2}$ | $3.6 \times 10^{-5}$ | -7.2 | -32.84 * |
| $r_{100}^2$ | $2.9650 \times 10^{-3}$ | $1.7 \times 10^{-5}$ | $3.1556 \times 10^{-3}$ | $1.8 \times 10^{-5}$ | -6.0 | -7.70 * |
| $r_{150}^2$ | $1.5374 \times 10^{-3}$ | $1.4 \times 10^{-5}$ | $1.6424 \times 10^{-3}$ | $1.6 \times 10^{-5}$ | -6.4 | -4.94 * |
| $\theta$ | $8.0025 \times 10^{-4}$ | $2.8 \times 10^{-7}$ | $7.9966 \times 10^{-4}$ | $3.4 \times 10^{-7}$ | 0.1 | 1.34 |
| S | $1.7923 \times 10^4$ | 5.0 | $1.7916 \times 10^4$ | 6.1 | 0.0 | 0.89 |
| $S_{<.1}$ | $9.5630 \times 10^3$ | 3.9 | $9.5639 \times 10^3$ | 4.4 | -0.0 | -0.15 |
| $S_{\in[.1,.4]}$ | $6.5711 \times 10^3$ | 3.0 | $6.5642 \times 10^3$ | 3.4 | 0.1 | 1.52 |
| $S_{>.4}$ | $1.7890 \times 10^3$ | 1.6 | $1.7879 \times 10^3$ | 1.8 | 0.1 | 0.46 |

**Table 46:**
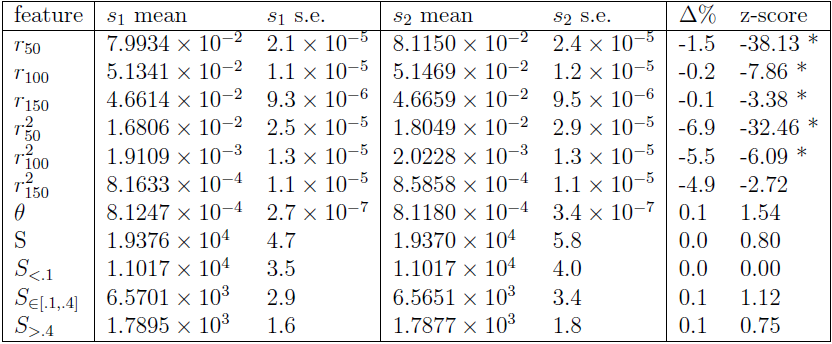
*s*_1_ = COSI2 with “-u” set to 0, *s*_2_ = MS; model: exponential expansion

| feature | $s_1$ mean | $s_1$ s.e. | $s_2$ mean | $s_2$ s.e. | $\Delta\%$ | z-score |
| --- | --- | --- | --- | --- | --- | --- |
| $r_{50}$ | $7.9934 \times 10^{-2}$ | $2.1 \times 10^{-5}$ | $8.1150 \times 10^{-2}$ | $2.4 \times 10^{-5}$ | -1.5 | -38.13 * |
| $r_{100}$ | $5.1341 \times 10^{-2}$ | $1.1 \times 10^{-5}$ | $5.1469 \times 10^{-2}$ | $1.2 \times 10^{-5}$ | -0.2 | -7.86 * |
| $r_{150}$ | $4.6614 \times 10^{-2}$ | $9.3 \times 10^{-6}$ | $4.6659 \times 10^{-2}$ | $9.5 \times 10^{-6}$ | -0.1 | -3.38 * |
| $r_{50}^2$ | $1.6806 \times 10^{-2}$ | $2.5 \times 10^{-5}$ | $1.8049 \times 10^{-2}$ | $2.9 \times 10^{-5}$ | -6.9 | -32.46 * |
| $r_{100}^2$ | $1.9109 \times 10^{-3}$ | $1.3 \times 10^{-5}$ | $2.0228 \times 10^{-3}$ | $1.3 \times 10^{-5}$ | -5.5 | -6.09 * |
| $r_{150}^2$ | $8.1633 \times 10^{-4}$ | $1.1 \times 10^{-5}$ | $8.5858 \times 10^{-4}$ | $1.1 \times 10^{-5}$ | -4.9 | -2.72 |
| $\theta$ | $8.1247 \times 10^{-4}$ | $2.7 \times 10^{-7}$ | $8.1180 \times 10^{-4}$ | $3.4 \times 10^{-7}$ | 0.1 | 1.54 |
| S | $1.9376 \times 10^4$ | 4.7 | $1.9370 \times 10^4$ | 5.8 | 0.0 | 0.80 |
| $S_{<.1}$ | $1.1017 \times 10^4$ | 3.5 | $1.1017 \times 10^4$ | 4.0 | 0.0 | 0.00 |
| $S_{\in[.1,.4]}$ | $6.5701 \times 10^3$ | 2.9 | $6.5651 \times 10^3$ | 3.4 | 0.1 | 1.12 |
| $S_{>.4}$ | $1.7895 \times 10^3$ | 1.6 | $1.7877 \times 10^3$ | 1.8 | 0.1 | 0.75 |

#### 3.11 ARGON and MS, with gene conversion

**Table 47:** *s*_1_ = ARGON (with gene conversion), *s*_2_ = MS (with gene conversion); model: constant size

| feature | $s_1$ mean | $s_1$ s.e. | $s_2$ mean | $s_2$ s.e. | $\Delta\%$ | z-score |
| --- | --- | --- | --- | --- | --- | --- |
| $r_{50}$ | $9.0238 \times 10^{-2}$ | $3.3 \times 10^{-5}$ | $9.0292 \times 10^{-2}$ | $3.4 \times 10^{-5}$ | -0.1 | -1.14 |
| $r_{100}$ | $5.8399 \times 10^{-2}$ | $2.1 \times 10^{-5}$ | $5.8396 \times 10^{-2}$ | $2.1 \times 10^{-5}$ | 0.0 | 0.10 |
| $r_{150}$ | $5.2513 \times 10^{-2}$ | $1.8 \times 10^{-5}$ | $5.2508 \times 10^{-2}$ | $1.9 \times 10^{-5}$ | 0.0 | 0.19 |
| $r_{50}^2$ | $2.0488 \times 10^{-2}$ | $3.3 \times 10^{-5}$ | $2.0541 \times 10^{-2}$ | $3.4 \times 10^{-5}$ | -0.3 | -1.12 |
| $r_{100}^2$ | $3.1214 \times 10^{-3}$ | $1.7 \times 10^{-5}$ | $3.1110 \times 10^{-3}$ | $1.7 \times 10^{-5}$ | 0.3 | 0.43 |
| $r_{150}^2$ | $1.6233 \times 10^{-3}$ | $1.4 \times 10^{-5}$ | $1.6329 \times 10^{-3}$ | $1.5 \times 10^{-5}$ | -0.6 | -0.47 |
| $\theta$ | $7.9951 \times 10^{-4}$ | $3.1 \times 10^{-7}$ | $8.0003 \times 10^{-4}$ | $3.3 \times 10^{-7}$ | -0.1 | -1.15 |
| S | $1.7911 \times 10^4$ | 5.7 | $1.7917 \times 10^4$ | 6.0 | -0.0 | -0.72 |
| $S_{<.1}$ | $9.5604 \times 10^3$ | 4.2 | $9.5592 \times 10^3$ | 4.4 | 0.0 | 0.20 |
| $S_{\in[.1,.4]}$ | $6.5622 \times 10^3$ | 3.2 | $6.5672 \times 10^3$ | 3.4 | -0.1 | -1.07 |
| $S_{>.4}$ | $1.7885 \times 10^3$ | 1.6 | $1.7903 \times 10^3$ | 1.7 | -0.1 | -0.77 |

**Table 48:** *s*_1_ = ARGON (with gene conversion), *s*_2_ = MS (with gene conversion); model: exponential expansion

| feature | $s_1$ mean | $s_1$ s.e. | $s_2$ mean | $s_2$ s.e. | $\Delta\%$ | z-score |
| --- | --- | --- | --- | --- | --- | --- |
| $r_{50}$ | $7.9468 \times 10^{-2}$ | $2.2 \times 10^{-5}$ | $7.9555 \times 10^{-2}$ | $2.3 \times 10^{-5}$ | -0.1 | -2.73 |
| $r_{100}$ | $5.1365 \times 10^{-2}$ | $1.1 \times 10^{-5}$ | $5.1389 \times 10^{-2}$ | $1.2 \times 10^{-5}$ | -0.0 | -1.47 |
| $r_{150}$ | $4.6628 \times 10^{-2}$ | $9.3 \times 10^{-6}$ | $4.6630 \times 10^{-2}$ | $9.4 \times 10^{-6}$ | -0.0 | -0.15 |
| $r_{50}^2$ | $1.6708 \times 10^{-2}$ | $2.6 \times 10^{-5}$ | $1.6756 \times 10^{-2}$ | $2.6 \times 10^{-5}$ | -0.3 | -1.31 |
| $r_{100}^2$ | $2.0119 \times 10^{-3}$ | $1.2 \times 10^{-5}$ | $2.0245 \times 10^{-3}$ | $1.2 \times 10^{-5}$ | -0.6 | -0.74 |
| $r_{150}^2$ | $8.5637 \times 10^{-4}$ | $1.1 \times 10^{-5}$ | $8.6228 \times 10^{-4}$ | $1.1 \times 10^{-5}$ | -0.7 | -0.38 |
| $\theta$ | $8.1226 \times 10^{-4}$ | $3.2 \times 10^{-7}$ | $8.1187 \times 10^{-4}$ | $3.3 \times 10^{-7}$ | 0.0 | 0.85 |
| S | $1.9372 \times 10^4$ | 5.4 | $1.9361 \times 10^4$ | 5.7 | 0.1 | 1.40 |
| $S_{<.1}$ | $1.1013 \times 10^4$ | 3.8 | $1.1007 \times 10^4$ | 4.0 | 0.1 | 1.09 |
| $S_{\in[.1,.4]}$ | $6.5697 \times 10^3$ | 3.3 | $6.5643 \times 10^3$ | 3.3 | 0.1 | 1.16 |
| $S_{>.4}$ | $1.7893 \times 10^3$ | 1.7 | $1.7900 \times 10^3$ | 1.7 | -0.0 | -0.29 |

#### 3.12 ARGON and COSI2, with gene conversion

**Table 49:** *s*_1_ = ARGON (with gene conversion), *s*_2_ = COSI2 (with gene conversion); model: constant size

| feature | $s_1$ mean | $s_1$ s.e. | $s_2$ mean | $s_2$ s.e. | $\Delta\%$ | z-score |
| --- | --- | --- | --- | --- | --- | --- |
| $r_{50}$ | $9.0238 \times 10^{-2}$ | $3.3 \times 10^{-5}$ | $9.0289 \times 10^{-2}$ | $3.4 \times 10^{-5}$ | -0.1 | -1.08 |
| $r_{100}$ | $5.8399 \times 10^{-2}$ | $2.1 \times 10^{-5}$ | $5.8429 \times 10^{-2}$ | $2.1 \times 10^{-5}$ | -0.1 | -1.01 |
| $r_{150}$ | $5.2513 \times 10^{-2}$ | $1.8 \times 10^{-5}$ | $5.2524 \times 10^{-2}$ | $1.8 \times 10^{-5}$ | -0.0 | -0.43 |
| $r_{50}^2$ | $2.0488 \times 10^{-2}$ | $3.3 \times 10^{-5}$ | $2.0469 \times 10^{-2}$ | $3.4 \times 10^{-5}$ | 0.1 | 0.40 |
| $r_{100}^2$ | $3.1214 \times 10^{-3}$ | $1.7 \times 10^{-5}$ | $3.1174 \times 10^{-3}$ | $1.7 \times 10^{-5}$ | 0.1 | 0.17 |
| $r_{150}^2$ | $1.6233 \times 10^{-3}$ | $1.4 \times 10^{-5}$ | $1.6433 \times 10^{-3}$ | $1.5 \times 10^{-5}$ | -1.2 | -0.97 |
| $\theta$ | $7.9951 \times 10^{-4}$ | $3.1 \times 10^{-7}$ | $7.9994 \times 10^{-4}$ | $3.3 \times 10^{-7}$ | -0.1 | -0.95 |
| S | $1.7911 \times 10^4$ | 5.7 | $1.7914 \times 10^4$ | 5.9 | -0.0 | -0.37 |
| $S_{<.1}$ | $9.5604 \times 10^3$ | 4.2 | $9.5560 \times 10^3$ | 4.3 | 0.0 | 0.73 |
| $S_{\in[.1,.4]}$ | $6.5622 \times 10^3$ | 3.2 | $6.5701 \times 10^3$ | 3.4 | -0.1 | -1.69 |
| $S_{>.4}$ | $1.7885 \times 10^3$ | 1.6 | $1.7874 \times 10^3$ | 1.7 | 0.1 | 0.47 |

**Table 50:** *s*_1_ = ARGON (with gene conversion), *s*_2_ = COSI2 (with gene conversion); model: exponential expansion

| feature | $s_1$ mean | $s_1$ s.e. | $s_2$ mean | $s_2$ s.e. | $\Delta\%$ | z-score |
| --- | --- | --- | --- | --- | --- | --- |
| $r_{50}$ | $7.9468 \times 10^{-2}$ | $2.2 \times 10^{-5}$ | $7.9472 \times 10^{-2}$ | $2.3 \times 10^{-5}$ | -0.0 | -0.13 |
| $r_{100}$ | $5.1365 \times 10^{-2}$ | $1.1 \times 10^{-5}$ | $5.1373 \times 10^{-2}$ | $1.2 \times 10^{-5}$ | -0.0 | -0.49 |
| $r_{150}$ | $4.6628 \times 10^{-2}$ | $9.3 \times 10^{-6}$ | $4.6625 \times 10^{-2}$ | $9.4 \times 10^{-6}$ | 0.0 | 0.23 |
| $r_{50}^2$ | $1.6708 \times 10^{-2}$ | $2.6 \times 10^{-5}$ | $1.6733 \times 10^{-2}$ | $2.6 \times 10^{-5}$ | -0.1 | -0.68 |
| $r_{100}^2$ | $2.0119 \times 10^{-3}$ | $1.2 \times 10^{-5}$ | $2.0191 \times 10^{-3}$ | $1.3 \times 10^{-5}$ | -0.4 | -0.41 |
| $r_{150}^2$ | $8.5637 \times 10^{-4}$ | $1.1 \times 10^{-5}$ | $8.7108 \times 10^{-4}$ | $1.1 \times 10^{-5}$ | -1.7 | -0.95 |
| $\theta$ | $8.1226 \times 10^{-4}$ | $3.2 \times 10^{-7}$ | $8.1193 \times 10^{-4}$ | $3.4 \times 10^{-7}$ | 0.0 | 0.71 |
| S | $1.9372 \times 10^4$ | 5.4 | $1.9364 \times 10^4$ | 5.8 | 0.0 | 1.01 |
| $S_{<.1}$ | $1.1013 \times 10^4$ | 3.8 | $1.1008 \times 10^4$ | 4.0 | 0.0 | 0.91 |
| $S_{\in[.1,.4]}$ | $6.5697 \times 10^3$ | 3.3 | $6.5684 \times 10^3$ | 3.4 | 0.0 | 0.27 |
| $S_{>.4}$ | $1.7893 \times 10^3$ | 1.7 | $1.7882 \times 10^3$ | 1.7 | 0.1 | 0.46 |

#### 3.13 MS and COSI2, with gene conversion

**Table 51:** *s*_1_ = MS (with gene conversion), *s*_2_ = COSI2 (with gene conversion); model: constant size

| feature | $s_1$ mean | $s_1$ s.e. | $s_2$ mean | $s_2$ s.e. | $\Delta\%$ | z-score |
| --- | --- | --- | --- | --- | --- | --- |
| $r_{50}$ | $9.0292 \times 10^{-2}$ | $3.4 \times 10^{-5}$ | $9.0289 \times 10^{-2}$ | $3.4 \times 10^{-5}$ | 0.0 | 0.06 |
| $r_{100}$ | $5.8396 \times 10^{-2}$ | $2.1 \times 10^{-5}$ | $5.8429 \times 10^{-2}$ | $2.1 \times 10^{-5}$ | -0.1 | -1.11 |
| $r_{150}$ | $5.2508 \times 10^{-2}$ | $1.9 \times 10^{-5}$ | $5.2524 \times 10^{-2}$ | $1.8 \times 10^{-5}$ | -0.0 | -0.61 |
| $r_{50}^2$ | $2.0541 \times 10^{-2}$ | $3.4 \times 10^{-5}$ | $2.0469 \times 10^{-2}$ | $3.4 \times 10^{-5}$ | 0.4 | 1.50 |
| $r_{100}^2$ | $3.1110 \times 10^{-3}$ | $1.7 \times 10^{-5}$ | $3.1174 \times 10^{-3}$ | $1.7 \times 10^{-5}$ | -0.2 | -0.27 |
| $r_{150}^2$ | $1.6329 \times 10^{-3}$ | $1.5 \times 10^{-5}$ | $1.6433 \times 10^{-3}$ | $1.5 \times 10^{-5}$ | -0.6 | -0.49 |
| $\theta$ | $8.0003 \times 10^{-4}$ | $3.3 \times 10^{-7}$ | $7.9994 \times 10^{-4}$ | $3.3 \times 10^{-7}$ | 0.0 | 0.19 |
| S | $1.7917 \times 10^4$ | 6.0 | $1.7914 \times 10^4$ | 5.9 | 0.0 | 0.36 |
| $S_{<.1}$ | $9.5592 \times 10^3$ | 4.4 | $9.5560 \times 10^3$ | 4.3 | 0.0 | 0.52 |
| $S_{\in[.1,.4]}$ | $6.5672 \times 10^3$ | 3.4 | $6.5701 \times 10^3$ | 3.4 | -0.0 | -0.60 |
| $S_{>.4}$ | $1.7903 \times 10^3$ | 1.7 | $1.7874 \times 10^3$ | 1.7 | 0.2 | 1.21 |

**Table 52:** *s*_1_ = MS (with gene conversion), *s*_2_ = COSI2 (with gene conversion); model: exponential expansion

| feature | $s_1$ mean | $s_1$ s.e. | $s_2$ mean | $s_2$ s.e. | $\Delta\%$ | z-score |
| --- | --- | --- | --- | --- | --- | --- |
| $r_{50}$ | $7.9555 \times 10^{-2}$ | $2.3 \times 10^{-5}$ | $7.9472 \times 10^{-2}$ | $2.3 \times 10^{-5}$ | 0.1 | 2.55 |
| $r_{100}$ | $5.1389 \times 10^{-2}$ | $1.2 \times 10^{-5}$ | $5.1373 \times 10^{-2}$ | $1.2 \times 10^{-5}$ | 0.0 | 0.94 |
| $r_{150}$ | $4.6630 \times 10^{-2}$ | $9.4 \times 10^{-6}$ | $4.6625 \times 10^{-2}$ | $9.4 \times 10^{-6}$ | 0.0 | 0.38 |
| $r_{50}^2$ | $1.6756 \times 10^{-2}$ | $2.6 \times 10^{-5}$ | $1.6733 \times 10^{-2}$ | $2.6 \times 10^{-5}$ | 0.1 | 0.63 |
| $r_{100}^2$ | $2.0245 \times 10^{-3}$ | $1.2 \times 10^{-5}$ | $2.0191 \times 10^{-3}$ | $1.3 \times 10^{-5}$ | 0.3 | 0.31 |
| $r_{150}^2$ | $8.6228 \times 10^{-4}$ | $1.1 \times 10^{-5}$ | $8.7108 \times 10^{-4}$ | $1.1 \times 10^{-5}$ | -1.0 | -0.57 |
| $\theta$ | $8.1187 \times 10^{-4}$ | $3.3 \times 10^{-7}$ | $8.1193 \times 10^{-4}$ | $3.4 \times 10^{-7}$ | -0.0 | -0.13 |
| S | $1.9361 \times 10^4$ | 5.7 | $1.9364 \times 10^4$ | 5.8 | -0.0 | -0.37 |
| $S_{<.1}$ | $1.1007 \times 10^4$ | 4.0 | $1.1008 \times 10^4$ | 4.0 | -0.0 | -0.18 |
| $S_{\in[.1,.4]}$ | $6.5643 \times 10^3$ | 3.3 | $6.5684 \times 10^3$ | 3.4 | -0.1 | -0.87 |
| $S_{>.4}$ | $1.7900 \times 10^3$ | 1.7 | $1.7882 \times 10^3$ | 1.7 | 0.1 | 0.75 |

